# Neuronal extracellular vesicles regulate axon development in primary cortical neurons via local miR-99a targeting of HS3ST2

**DOI:** 10.1101/2025.10.24.684188

**Authors:** Raquel Mesquita-Ribeiro, Alex Rathbone, Paula Jackson, Sophie McCann, Hannah Jackson, Cristiano Lucci, Victoria James, Federico Dajas-Bailador

## Abstract

Fully functional neural competence and integrity requires a complex array of communication means among neurons, with extracellular vesicles (EVs) emerging as a relevant mechanism for cell-cell interaction in the CNS. Despite the growing number of studies demonstrating the presence of microRNAs (miRNAs) in axon and EVs, the molecular mechanisms of those miRNAs present in EVs and their functional role in nervous system development has not been fully explored. In this study, we investigated whether neuronal EVs can have a role in neuron-to-neuron communication during the development of neuron connectivity in mouse primary cortical neuron cultures. Our results demonstrate how miR-99a can regulate axonal growth via its EV-mediated delivery and through the targeting of HS3ST2, a heparan sulphate glucosamine 3-O-sulphotransferase, which is predominantly expressed in the brain and generates rare 3-O-sulphated domains in heparan sulphate proteoglycans, with growing importance in development and neurodegenerative mechanisms. Importantly, we show how in compartmentalised microfluidic cultures, where axons are isolated from neuronal somas, the growth-promoting effects of neuron-derived EVs are local to the axon. These findings establish that neuronal EVs can deliver miRNAs to discrete subcellular domains to acutely modulate local gene expression, thereby driving axonal growth and shaping neurodevelopment.

## INTRODUCTION

Neurons are highly complex and specialised cells, capable of conveying information over large distances by typically possessing several dendrites and a single long connective process, the axon. This complex and highly compartmentalised cellular morphology must in turn employ unique solutions to maintain and modify the local proteome that is required for the nervous system’s correct wiring during development, and for its function and plasticity during adulthood (1, 2). In effect, the precise sub-cellular localisation of messenger RNA, together with on-site protein synthesis, is a conserved mechanism that appears to have evolved to fulfil local demand at short time scales and to provide sub-cellular capabilities for neuronal development, synaptogenesis, experience-dependent plasticity, and survival (3).

In this context, fully functional neural competence and integrity requires a complex array of communication mechanisms among different types of neurons, such as transcellular signalling *via* synapses, the plasticity-inducing influence of specific patterns of electrical activity, and mechanisms of anterograde and retrograde axonal transport through neurotrophins, RNAs, growth factors and multiple signalling proteins and transcription factors (3–9). More recently, studies have also revealed an expanded range of modes of communication among cells, which includes extracellular vesicles (EVs), such as exosomes and microvesicles, and which has added another relevant layer of complexity to cell-cell interactions within the nervous system (4, 10).

EVs are produced by virtually all cells, including those in the central nervous system (CNS) (11), and as small membranous vesicles bounded by a lipid bilayer and secreted into the extracellular space, they can carry diverse intraluminal cargos such as proteins, lipids, and nucleic acids (12). They can be subdivided into microvesicles (100 nm to 1 μm in diameter) and exosomes (30 to 150 nm in diameter), depending on their subcellular origin from plasma membrane or endocytic pathway, respectively (13). Even though the study of EVs is in its early stages, they are known to play vital roles in the intercellular communication that underlies various physiological processes and pathological functions of both recipient and parent cells (12, 14). In cancer models, the role of EV-mediated signalling has been linked to various pathological processes, from promotion of premetastatic niche in target tissues (15), to autocrine processes that accelerate tumour growth progression (16). However, besides early evidence that neuronal cultures can regulate EV release (17–21), and the more recent demonstration for neuron-derived EV mediated function in neurogenesis (22), and inter-neuronal exchange and neurotransmission regulation via vesicular synaptobrevin (23), their role in nervous system development has only begun to be unravelled,

Interest in EVs was significantly stoked by the observation that they can contain nucleic acids and mediate the transfer of RNAs between cells (24, 25), including those of the nervous system (26, 27). Various types of RNAs have been detected within EVs, such as circular RNA, ribosomal RNA, and many other small noncoding RNAs (28–30). Indeed, our recent work highlighted the highly specific content of small non-coding RNAs (sncRNAs) in neuronal axons and EVs derived from mouse primary cortical neurons, which included microRNAs but also the precisely processed fragments derived from tRNAs and snRNAs (31).

In the context of post transcriptional regulation of cell function, miRNAs stand out for their capacity for reprogramming protein expression in localised subcellular domains and have emerged as important players in multiple cellular processes, such as neurogenesis, axon development, pathfinding and neuron connectivity (32–36). Despite the growing number of studies demonstrating the presence of miRNAs in axon and EVs (31, 37), the molecular mechanisms of miRNAs in cell-cell communication and its functional effects at the local level in the nervous system has only begun to be explored. Indeed, although changes in miRNA levels have been traditionally regarded as cell intrinsic, there is now mounting evidence for their cell transfer (38) and even purely extracellular roles (39). Among these growing number of processes, one of the key cellular mechanisms that enables sncRNAs to be transferred between cells is their selective packaging into extracellular vesicles (EVs) and trafficking in the extracellular space.

As one of the many miRNAs known to have a role in the nervous system (40–42), miR-99a ranked within the top 20 highly expressed miRNAs over development in rat cerebral cortex up to 28 days postnatal (43). Furthermore, miR-99a was also found upregulated in a microRNA expression profile during the transition from neuronal progenitors into the earliest differentiating rat cortical neurons (40). Consistent with a role for miR-99a in neuron subcellular compartments, both its mature and precursor transcripts were found to be enriched in synaptic fractions of mouse forebrain (44) can be secreted from synaptosomes via depolarisation and Ca^2+^-dependent mechanisms (45). More recently, work in our lab identified miR-99a as part of a subset of 23 conserved miRNAs detected across axonal samples in different neuronal subtypes and species, and across in vitro and in vivo models (31). In this same study, miR-99a was among the most highly expressed miRNAs in neuron-derived EVs. Despite this growing amount of information, evidence of a precise functional mechanism for miR-99a in the nervous system is still scarce.

The capacity for miRNA transfer via EVs in cells from the nervous system has been previously shown in astrocytes, which can transfer miRNAs to metastatic tumour cells, thus promoting brain metastasis (46). Importantly, the miRNA cargo of EVs produced by glial cells may also regulate the expression of neuronal genes (47, 48). In effect, although glial to neuron communication has been the centre of most studies on EV mediated transfer in the nervous system, neuron-to-neuron communication has also been detected (10, 22, 23, 49), including recent demonstration that EVs derived from cortical cultures modulate synaptic activity of recipient hippocampal cultures (50) and neuron-derived EVs promote spine formation and preserve neuronal complexity in primary cortical cultures (51). Moreover, neuronal EVs have been show to mediate the inter-neuronal delivery of miRNAs able to induce BDNF-dependent changes in circuit connectivity (52), while miR-21-5p has been shown to be dynamically released in response to hypoxia in primary cortical neurons (53) and to functionally mediate sensory neuron to macrophage EV communication after nerve trauma (54, 55), which can influence the nature of immune infiltrate in the dorsal root ganglia and noxious signalling. Overall, these findings have started to unravel the potential for neuron-derived EVs in the miRNA-mediated modulation of neuronal systems and networks.

In this study, we investigated whether EVs can have a role in neuron communication during the development of neuronal networks *in vitro*. Our results show how miR-99a can regulate axonal growth via its EV-mediated delivery and through the targeting of HS3ST2, a heparan sulphate glucosamine 3-O-sulphotransferase, which is predominantly expressed in the brain and generates rare 3-O-sulphated domains in heparan sulphate proteoglycans, with growing importance in vertebrate development (56) and neurodegenerative mechanisms (57).

## MATERIALS AND METHODS

### Primary cortical neuron cultures

Mice (C57/BL6) were housed and bred in compliance with the ethics of animal welfare and in accordance to the *Animal* (*Scientific Procedures*) *Act 1986.* C57/BL6 mouse embryos at E16.5 stage of development were culled, and their brains removed. Brain cortices were dissected, and the meninges separated under a dissection microscope. The tissue was further incubated in Hanks Balanced Salt Solution (HBSS, Ca^2+^ and Mg^2+^-free; Gibco) with 1 mg/ml trypsin and 5 mg/ml DNAse I (Sigma-Aldrich) at 37°C for 30’. Following the addition of 0.05% (v/v) soybean trypsin inhibitor (Sigma-Aldrich), the tissue was mechanically dissociated in Neurobasal media (Invitrogen) supplemented with 1X GlutaMax and 2% B-27 (Gibco). Dissociated neurons were resuspended in supplemented Neurobasal media (10×10^6^ cells/mL). For neuron transfections and RNA extraction, neurons were plated at a final seeding density of 1.75×10^5^ cells/cm^2^ in 6-well plates (Corning) with or without 22×22mm glass coverslips (Menzel Glaser) and previously coated with 50 µg/ml poly-L-ornithine (PLO; Sigma-Aldrich). For immunostainings and functional assays with EVs, neurons were plated at a final density of 3.5×10^3^/mL in 12-well plates (Corning) with PLO-coated 19mm glass coverslips (Menzel Glaser). For experiments that required over 7 days in culture, media was replenished with ¼ of its volume every 2-3 days.

### EV isolation

The conditioned media of primary cortical neurons cultured for 2 weeks *in vitro* in four 6-well plates (seeding density 1.75 × 105 cells/cm2) was collected per individual preparation. Pooled culture media (∼48 mL) underwent filtration using a 10 K MWCO centrifugal concentrator (PES, 20mL concentrator, Pierce) for 30 min/4000 g to a final volume of 500uL. Extracellular vesicle (EV) fractions were further isolated by size-exclusion chromatography (SEC) using commercially available sepharose columns (pore 70 nm, qEV IZON), in which the EV fraction in the media is separated by gravity flow (Fig. 1.D), using Ca2+/Mg2+-free PBS as elution buffer, in accordance with manufacturer’s procedure to isolate small EVs <200 nm and within the range considered to include exosomes (58). All three of the EV fractions were pooled together for subsequent experiments. This method attempts minimal disturbance of vesicle size and content by avoiding ultracentifugation forces, thus best preserving EV integrity and, importantly, their biological activity (59).

**Figure 1.**
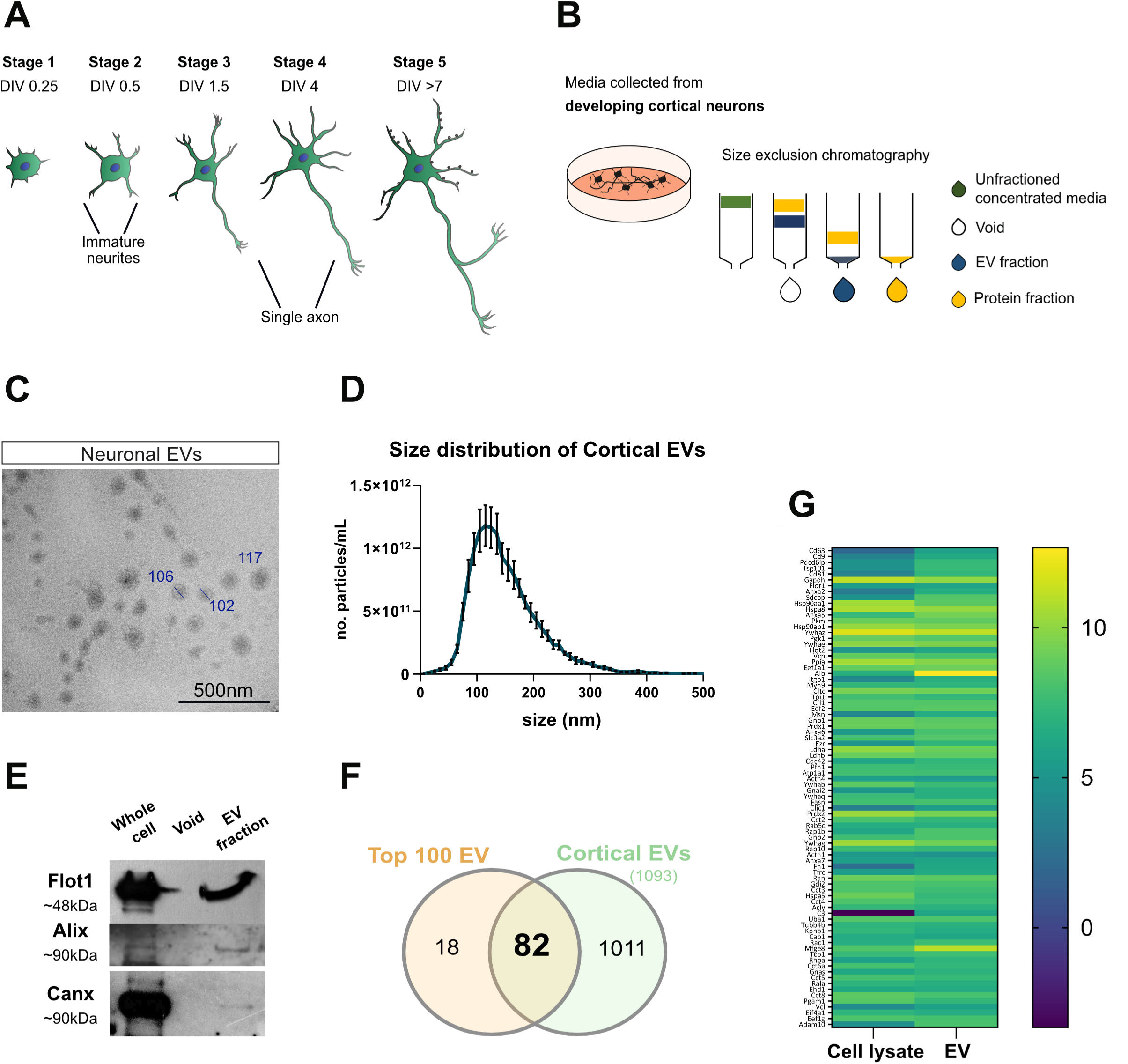
Mouse primary cortical neurons generate extracellular vesicles (EVs) in culture. **(A)** Schematic representation for the development of mouse primary cortical neurons in culture, depicting axon specification within the first 24h after seeding (stage 2-3) and early axon elongation during the first week (stage 3-4), followed by synaptogenesis after the first week in culture (stage 5). This study investigated the period of intense axon outgrowth between DIV2-3 and DIV5. **(B)** Neuron-derived EVs were isolated from the conditioned media of primary cortical neuron cultures at 2 weeks. Diagrammatic representation of the size exclusion chromatography method used to separate the EV-enriched fractions from the free protein fraction present in the media (sepharose pore 70nm, qEV system) after an ultrafiltration step to concentrate the media (10K MWCO, 4000g/30min). **(C)** Electron microscopy imaging showed membrane-bound particles with the cup-shape morphology, characteristic of EVs (negative stain, scale bar:500nm). **(D)** Size distribution of cortical EV-enriched fractions by nanoparticle tracking analysis show a higher proportion of particles in the 50-200nm diameter range, with an average particle size of 137.7nm and average density of 2.1×10^12^ particles/mL across 3 independent EV isolations. **(E)** EV cargo characterization by immunoblotting showed the presence of EV markers Flotilin-1 and Alix and absence of ER marker Calnexin in EV fractions. **(F)** Proteomic profiling using SWATH mass spectrometry showed 82 of the top 100 EV-defining proteins were present in cortical EV samples. **(G)** A higher abundance of the top EV-defining proteins were present in the cortical EV samples compared to whole-cell lysate samples.

### EV characterisation

Following EV isolation, the EV fraction was analysed by nanoparticle tracking analysis (NTA), a method that can visualize and measure nanoparticles in suspension (range 10-1000 nm) and is based on the analysis of Brownian motion. NTA was performed as described in Mesquita-Ribeiro et al. (31) with minor modifications, using the PMX 220 TWIN ZetaView (Particle Metrix, Meerbusch, Germany). The parameters for all NTA measurements were as follows: sensitivity 80, shutter speed 100, with a frame rate of 30 frames per second. For all samples 2 technical assessments were carried out. For EV immunoblot analysis, PBS-washed cortical primary neurons and isolated EVs were lysed directly in gel loading buffer (0.15 M Tris, 8 M Urea, 2.5% SDS, 20% glycerol, 10% 2-mercaptoethanol, 3% DTT, 0.1% bromophenol blue) and loaded onto a 12% SDS-PAGE gel. Western blots were performed as described in Lucci et al., (36), using 10 ug total protein of whole cell lysate and 30 uL per SEC fraction (protein concentration not detectable by BCA protein assay kit, Pierce) and blotted for flotilin-1 (Santa Cruz; (C7) sc-133, 153, 1:1000) and Alix (Abcam: ab88388, 1:1000) as EV protein markers, and anti-calnexin (SicGen: AB0041-200, 1:1000 dilution), which is depleted in small EVs (58), as a negative control (Table 1). For transmission electron microscopy, EV samples were fixed in 3% glutaraldehyde in 0.1 M cacodylate buffer. 10 µls of fixed EVs were loaded onto poly-L-lysine coated copper grids and left to settle for 15 minutes. The excess liquid was wicked away with filter paper and the grids washed twice in Milli-Q water for 30 seconds, removing the excess water after each wash. The samples were stained with 10 µls of 2% Uranyl acetate (0.2 µm filtered) for 1 minute after which, excess stain was wicked away using filter paper and the grids was left to air dry. Grids were visualized using a Tecnai G2 T12 Biotwin transmission electron microscope (FEI) with an accelerating voltage of 100 kV and images of varying magnifications were captured using a MegaView II (Olympus) camera system.

### Proteomic analysis by SWATH-MS

Cell lysate samples (50 μg, n=3) were prepared in RIPA buffer following BCA quantification. EV samples were pooled and concentrated from all EV-enriched fractions (n=3). All samples processing and LC-MS/MS and SWATH analysis were performed at Nottingham Trent University. Proteins were digested using S-trap™ Micro spin columns. Samples were first dried at 60 °C and resuspended in 5% SDS/100 mM TEAB (pH 7.55). Samples were then reduced with 1 μl 0.5M DTT (20 min, 56 °C with shaking) and alkylated with 2 μl 0.5M iodoacetamide (15 min, RT, dark). Acidification with *2.8μl* 12% phosphoric acid (1:10 v/v) preceded protein trapping.

Each sample received 185 μl S-Trap buffer (90% methanol/100 mM TEAB, pH 7.1) before loading onto the S-trap (Protifi) micro spin columns and centrifuged (4000 × g, 30s). After four washes with 150 μl S-Trap buffer, the columns were transferred for digestion. Proteins were digested in 25 μl trypsin/EDTA (1 trypsin:10 protein (m/m)) in 50 mM TEAB (pH 7.5–8.0) for 1.5 h at 47 °C. Peptides were sequentially eluted with 50 mM TEAB, 0.2% formic acid, and 50% acetonitrile/0.2% formic acid, pooled, and dried at 60 °C. Dried peptides were resuspended in 5% acetonitrile/0.2% formic acid (30 μl for EVs and lysates), and 2 μl were injected by auto-sampler (Waters M-class) and gradient elution at 10 µL/min (Phenomenex Kinetex XB-C18 2.6 µm 15 x 0.3 mm analytical column, 30°C, mobile phase A 0.1% formic acid:mobile phase B acetonitrile with 0.1% formic acid LCMS grade) onto a Sciex ZenoTOF 7600. Gradient elution with the following profile: 3% B to 35% over 12 min, 80% B at 13-15 min, and re-equilibration at 12 µL/min to the starting conditions at 15.5 min for a 16.5 min total runtime. MS analysis was done via the Optiflow source with a micro flow 1-50 µL electrode in positive SWATH mode with zeno pulsing enabled at 4500 V, using 65 variable SWATH windows optimized on a complex human protein lysate from m/z 400-750 12 ms per window following a TOFMS scan of 25 ms for a total cycle time of 1.146 s. Ionization was deactivated at 14.2 min to reduce contamination during the column wash. SWATH data (.wiff format) was processed using DIA-NN 1.8.1 using the FASTA digest for library-free and Deep learning-based spectra options (Swissprot Human May 2022 with addition of TP53 isoforms and the cRAP proteome for contaminants) with the following parameters changed from default: FAST Digest for library-free search and Deep learning on, 1 missed cleavage, max variable mods 1 with Ox(M) selected, heuristic protein inference off. SWATH datasets were normalised by overlaying total ion traces and standardising to 5 μg input. Relative protein abundance across samples was quantified, while peptide hits shared by multiple proteins were excluded to avoid ambiguity. The top 100 EV proteins in Vesiclepedia database detected in at least 2 out of 3 biological replicates were considered.

### EV transfer

For the demonstration of neuron-neuron EV transfer, isolated EVs were incubated with BioTracker^TM^ MemBright 560 Live Cell Dye (SCT084, Sigma-Aldrich) at 200 nM in PBS (final concentration) for 30 minutes at room temperature in the dark. Following incubation, 50 µL PBS was then added to each sample. To remove unbound excess dye EVs were passed through Vesi-SEC micro size exclusion chromatography spin-columns (Vesiculab, Nottingham, UK) and used immediately after. As controls, PBS alone or mock EV fractions from fresh media were processed similarly to EVs, labelled with MemBright and cleared using Vesi-SEC columns. Next, 50 µL of cleared EVs (1×10^12^ particles/ml) and controls were added to cortical neuron cultures before imaging at varying timepoints using a Thermo Fisher EVOS M7000 Imaging System, with a 20X objective.

### Neuron functional assays

Neuronal transfections were performed 24 h after plating using 5 µL/well of Lipofectamine 2000 reagent and 250 µl/well of Opti-MEM reduced serum media (Thermo Fisher Scientific), in accordance with manufacturer’s instructions, and fixed 72 h after transfection. miRCURY LNA (Locked Nucleic Acid) microRNA inhibitor [50 nM], inhibitor control [50 nM], mimic [50 nM] and mimic control [50 nM], GapmeR control [50 nM], and GapmeR [50 nM] of miR-99a-5p (all Qiagen), referred to as miR-99a throughout this manuscript for simplicity, were used for miRNA functional assays. In all cases, 1 µg pmaxFP-Green (Lonza; hereafter referred to as GFP) was added to transfection mix for visualisation of transfected neurons. For protein overexpression studies, neurons were co-transfected with 1 µg GFP and either 1 µg of empty vector or 1 µg of pcDNA-HS3ST2.

In silencing experiments, a pool of siRNA specifically targeting *Hs3st2*, si*Hs3st2* (mouse; siGENOME SMARTpool) or siRNA non-targeting control (mouse; siGENOME SMARTpool #2) were co-transfected at final concentrations of 50 nM with 1ug GFP as reporter. Efficiency of siRNA Hs3st2 knockdown in primary cortical cultures was confirmed by qPCR, showing an average of ∼60% decrease in mRNA levels compared to non-targeting controls. For overexpression experiments 1 μg of pcDNA-HS3ST2 or empty vector were co-transfected with 1μg GFP.

To rescue the effects of miR-99a inhibition, cortical neurons were co-transfected with 1μg GFP and 25 nM inhibitor control or miR-99a inhibitor 25nM plus 35 nM siRNA control or si*Hs3st2*. To test the rescue of miR-99a ectopic expression, cortical cultures were co-transfected with 1ug GFP and either 30 nM mimic control or miR-99a mimic plus 1 µg of empty vector or 1 µg of pcDNA-HS3ST2.

In EV functional assays, ∼ 2.0×10^11^ EV particles in DPBS were added per condition, with or without miR-99a power inhibitor 100nM, at day in vitro 3 and incubated for 48h prior to fixation. The correspondent volume of DPBS was used for control conditions. To discard the possibility that media supplement contaminants could generate exogenous effects when co-purified with EVs, fresh culture media was used for “mock” EV isolation and corresponding fractions were tested for effects on axon development, as done previously. For experiments of miR-99a inhibition within EVs, the isolated EV fraction was incubated with miR-99a power inhibitor 100nM for 4h/RT, followed by SEC to isolate EVs from excess free miR-99a inhibitor.

In all experiments, cortical neurons were fixed in 4% paraformaldehyde (Thermo Fisher) and washed in PBS before direct visualisation and/or immunostaining. Microscope imaging was done using a widefield fluorescence microscope (Axiovert 200M, Zeiss), coupled to a CCD camera (Photometrics CoolSnap MYO) and Micro-Manager software 1.4.21 (60).

### RNAse, Proteinase K, Triton X-100 and dynasore assays

EV samples (∼ 2.0×10^11^ EV particles) in DPBS or equivalent buffer volume were incubated with or without 100µg/mL RNAse A (Thermo Fisher Scientific) or 50 μg/mL Proteinase K (AM2542, Ambion, Texas), and incubated for 30min at 37°C. RNAse A and proteinase K were neutralised with murine RNAse inhibitor (RiboLock Thermo Scientific) or mammalian proteinase inhibitor cocktail (1:100 Sigma), respectively. For detergent disruption assays, EV samples (∼ 2.0×10^11^ EV particles) in DPBS or equivalent buffer volume were treated with 0.5% and 0.075% Triton X-100, vortexed at maximum speed for 30sec and incubated at room temperature for 30min. The Triton-X concentrations used were shown to significantly disrupt EVs correspondent to exosome and microvesicle subpopulations (61). Treated EV fractions and DPBS controls were then loaded to DIV4 cortical neurons (seeding density 3.5×10^3^/mL) and fixed 6h later. For image analysis, neurons were immunolabelled for acetylated tubulin and axons were imaged and measured as described above. Efficiency of RNAse incubation was tested by RNA electrophoresis. In brief, 25ng of RNA isolated from primary cortical neurons was incubated with RNAse A (37°C/30min) and samples ran on High Sensitivity RNA ScreenTape, using the 4200 TapeStation System (Agilent; SuplFig2A). To test proteinase K activity, increasing amounts of BSA protein (2-20 ng) were incubated with proteinase K (37°C/30min), ran by SDS-PAGE and stained by Coomassie staining to visualise protein degradation. To test EV uptake in target neurons, dynasore (40 μM) was added for 30 min before addition of EVs for a further 6 hours. At the end of the incubation cells were fixed and axons measured as described previously.

### Compartmentalised neuronal cultures in microfluidic chambers

Primary cortical neurons were cultured for 6 days in microfluidic devices with 150 µm long microgrooves (SND150; Xona Microfluidics), a device that allows the fluidic isolation and functional compartmentalisation of the axon and somatodendritic compartments. For simplicity the somatodendritic compartment is hereafter designated the somal channel throughout the text. The devices were prepared as described previously (62). Briefly, ethanol sterile devices were mounted onto PLO-coated 35mm culture dishes (Nunc, ThermoFisher) and both channels equilibrated for 1 h with supplemented Neurobasal media. Following collection of excess media from the devices’ reservoirs, cortical neurons were added onto to the designated somatodendritic compartment at a seeding density of 4×10^6^ cells/ml and incubated for 30’ (37°C 5% CO_2_) to allow for cell attachment. The devices’ reservoirs were then topped up with supplemented Neurobasal media and cultures incubated at 37°C, 5% CO_2_. Axons were allowed to extend and cross the microgrooves to the axonal channel. EVs (∼5×10^10^ particles) were loaded to either the soma or axon compartments at 6 days *in vitro* and incubated for 48h. A difference in volume of ∼100 µl was always maintained between the somal and axonal channels in order to maintain fluidic isolation.

### Plasmid constructs and luciferase assay

For the pcDNA-HS3ST2 construct, *Hs3st2* cDNA was PCR-amplified from a replication construct (pGEM-HS3ST2; Stratech, Table 1) and the amplicon cloned into pcDNA3.1/Zeo(+) vector (kind gift from Dr Simon Dawson, University of Nottingham), using NheI/XbaI restriction sites.

For luciferase assays, a reporter construct containing the miR-99a-5p-binding site within the Hs3st2 3’UTR (*Hs3st2*) was cloned using using XhoI and NheI restriction sites in pmiRGLO dual reporter vector (Promega). Mutation of 3′ -UTR in miR-99a binding site of *Hs3st2* (*Hs3st2-*mut) was performed by site-directed mutagenesis of pmiRGLO-*Hs3st2* using primer pairs bearing a mutated seed sequence (mismatch at positions 2-5). All oligonucleotides used are listed in Table 1. HEK-293T cells were seeded in 24-well plates (Greiner) at a seeding density of 1.0×10^4^ cells/mL. After 24h, co-transfection of 100ng/well of pmirGLO-*Hs3st2* or *Hs3st2-mut* luciferase reporter construct and either miRNA mimic or mimic negative control at a final concentration of 100nM was performed in triplicate, using 1μL of Lipofectamine 2000 reagent and 100μL of Opti-MEM reduced serum media per well. Cells were harvested 48h after transfection in passive lysis buffer (Promega) for 30mins and further read with Dual Luciferase assay system (Promega) in the GloMax Multidetection System (Promega), following the manufacturer’s instructions. Each sample was read in duplicate and the firefly luciferase/renilla expression ratios were obtained. A total of three independent experiments were conducted with three technical replicates per condition. Per experiment, firefly luciferase/renilla expression ratios from each condition were normalised to respective control.

### RNA extraction and RT-qPCR

Cortical neurons grown on 6-well plates were scraped into 250 µL/well of TRIzol® Reagent (Fisher Scientific) and total RNA was isolated following manufacturer’s instructions and resuspended in RNAse-free water (Fisher Scientific) before storage at –80°C. For miRNA expression studies, cDNA was synthesised from mature miRNAs using the miRCURY LNA™ Universal cDNA synthesis kit (Qiagen, UK) as per manufacturer’s instructions, using 10 ng of total RNA. For each timepoint 5 biological samples were run in duplicate using miRCURY LNA™ primers (Table 1, Qiagen, UK). RT-qPCR was undertaken using the ExiLENT SYBR® Green master mix kit (Qiagen, UK). For mRNA targets, cDNA was synthesised from 100 ng total RNA (5 biological replicates), using SuperScript IV™ and Oligo(dT)_20_ primer (Invitrogen) as per manufacturer’s instructions. qPCR was undertaken using the PowerUp™ SYBR™ Green (Applied Biosystems) using 1.5 µL cDNA per replicate and 400 nM primers (Table 1, IDT). In both cases, PCR amplification was carried out in the Applied Biosystems Step One Plus thermocycler, using cycling parameters recommended by Qiagen (miRNA) and Applied Biosystems (mRNA). Data was acquired with Applied Biosystems SDS2.3 software. Passive reference dye ROX™ (Fisher Scientific) was included in all reactions. Expression data was analysed by relative quantification using the comparative Ct method (2^-ΔΔCt^). miR-99a-5p levels were analysed as relative expression to 4 h, using the geometric mean of miR-100-5p, miR-128-3p, miR-134-5p, miR-434-3p and let7a-5p as reference. miRNA reference genes were selected according to two parameters: detectable expression by RT-qPCR in axonal RNA samples and stable expression in previous in-house RT-qPCR studies on the development of cortical neuronal cultures (36). *Hs3st2 transcript* levels were analysed as relative expression to 4 h, where the geometric mean of *Gapdh* and *Ube2* levels was used as reference. All data are expressed as fold change to 4 h +/-SEM.

### Immunofluorescence

Cortical neurons cultured on coverslips or microfluidic devices were fixed using 4% paraformaldehyde (w/v) (ThermoFisher) for 30’, washed with 10 mM Glycine in PBS, permeabilised in PBS/Glycine-Triton (1x PBS, 10 mM Glycine, 0.2% Triton X-100; Sigma), blocked with 3% bovine serum albumin in PBS (BSA; Sigma) and further incubated with the appropriate primary antibodies overnight (Table 1). Following PBS-Triton 0.1% washes, cells were incubated with secondary antibodies (Alexa Fluor 488 and 568; 1:300 Molecular Probes) and mounted with Vectashield Hardset mounting media with Dapi (Vectorlabs).

### Data analysis

#### Measurement of axons in dissociated cortical cultures

For quantification of axon length, an axon was defined as a neurite that was at least 3 times the length of any other neurite and measured from the soma to the distal extent of the central region of the growth cone using Fiji software (63). Data are expressed as percentages of respective controls (∼300 axons measured for each condition from 4-6 independent experiments), mean ± SEM. The average length of axons in control groups was 115 ± 1.96 μm (mean ± SEM). In EV experiments, cultures were immunostained for acetylated tubulin and ∼100 neurons from ∼20 random fields were analysed per condition per experiment (∼400 axons from 4 independent experiments).

#### Quantification of fluorescence signal

Neurons labelled for HS3ST2 were imaged at 20x or 63x and images were further processed with Fiji software. Somas were manually selected and the area, mean grey value and integrated density were measured. In order to correct for background in each image, 3 empty areas were selected around every soma. Total cell fluorescence (C.F.) per cell was calculated as the measured integrated density corrected for background, according to the formula: C.F.= Integ.Density – [Area of soma X Average (mean grey value of background)]. For quantification of endogenous HS3ST2 over development in culture, a minimum of 20 images were randomly acquired and >60 neurons measured in each timepoint, DIV 2, 5 and 12. In each experiment, data from all timepoints was normalised to the average of C.F. at DIV2 and a total of three independent experiments were conducted.

For quantification of HS3ST2 levels after miR-99a mimic transfection, >50 GFP-positive neurons were imaged per condition and HS3ST2 C.F. analysed as above and normalised to the average C.F. of mimic control (3 independent experiments)

#### Measurement of axon length in microfluidic cortical cultures

The length of the axons was measured in Fiji software by tracing at least 125 axons in each condition from 3 independent experiments; each axon was traced from the edge of microgrooves to the growth cone of the longest axonal branch Data expressed as a percent of control (mean ± SEM).

### Statistical analysis

In all statistical tests, “n” refers to the number of independent experimental repeats, which varied from 4-8 depending on experimental model (see specific section for details). Data analysis was done using Prism v9.0 (GraphPad Software) and all data groups shown are expressed as mean ± SEM. The probability distribution of the data set was analysed before further statistical analysis (Shapiro–Wilk test). Statistical evaluation between two groups was performed using unpaired Student’s *t*-test. Analysis of more than 2 groups were carried out using 1-way ANOVA with Bonferroni post hoc analysis. Kruskal-Wallis’ test followed by a Dunn’s multiple comparisons test was used for non-parametric distributions. For all tests, * *p <* 0.05 and ** *p <* 0.01 were used as thresholds for statistical significant difference. For all tests P values are two-tailed.

## RESULTS

### Isolation and characterization of EVs from cortical primary neurons

To investigate the role of EV-mediated inter-neuronal communication in the development of CNS neurons *in vitro*, EVs were purified by size exclusion chromatography (SEC) from pooled media of primary cortical neurons cultured for 9-10 days (**Fig 1A-B**). Before functional studies, phase-contrast transmission electron microscopy confirmed the presence of a varied particle population, including typical bilayer-membrane vesicles that were heterogeneous in size, ranging in diameter from 50 to 120 nm (**Fig 1C**). Further nano particle tracking analysis showed EV particle density and size distribution in agreement with previous studies (**Fig 1D**) (17, 18, 31, 64).

As further test of the identity of this EV fraction we assessed specific markers by western blot and compared them side-by-side to the eluted protein fraction and parental whole cell lysate. As expected, expression of known EV markers flotillin-1 and alix was observed in the EV samples, while calnexin, an ER protein unlikely to be present in small EVs (58), was largely absent in our EV fractions when compared to whole cell samples (**Fig 1E**). At the same time, we used fluorescence microscopy to demonstrate the presence of flotillin-1 positive structures (65, 66) at early (DIV 2-3) and late (DIV 9) stages of development in primary neuron culture, DIV 2 (**Suppl Fig 1A-B**), including characteristic vesicular staining within cell bodies and axon growth cones (**Suppl Fig 1C**). Although flotilins are commonly included among canonical EV markers, they are ubiquitous membrane-associated proteins also present in endocytic and protein trafficking structures. Therefore, to obtain a more accurate characterisation of our purified EV fraction, we performed comprehensive proteomic profiling using SWATH mass spectrometry. As shown in **Figure 1F**, 82 of the top 100 EV-defining proteins, (Vesiclepedia, 2024; (67) were present in cortical EV samples (**Fig 1F**), and had higher abundance compared to whole-cell lysate samples (**Fig 1G**). Of note, Tsg101, tetraspanins CD63, CD9, CD81 and Flot1 are all among the most abundant proteins in our EV samples.

### Addition of neuron-derived EVs promotes axonal growth in primary cortical neurons

The abundance of flotillin-1 staining in somatic peri-nuclear areas, and its high concentration in dynamic cytoskeleton domains such as axon growth cones (**Suppl Fig 1**) prompted us to test whether EVs can display a functionally bioactive role in developing neurons. For this, we exposed early-stage cortical neurons (Day 3) to increasing amounts of EVs that have been previously isolated from the conditioned media of later-stage cultures (Day 9), whereas the same volume of the EV vehicle DPBS was used as a negative control (**Fig 2A**). Addition of increasing quantities of EVs to the 3 DIV neuronal cultures generated a concentration-dependent increase in axon outgrowth after 48 h (**Fig 2B-C**), suggesting a trophic and autocrine-like intrinsic function of the EV fraction in axon development.

**Figure 2.**
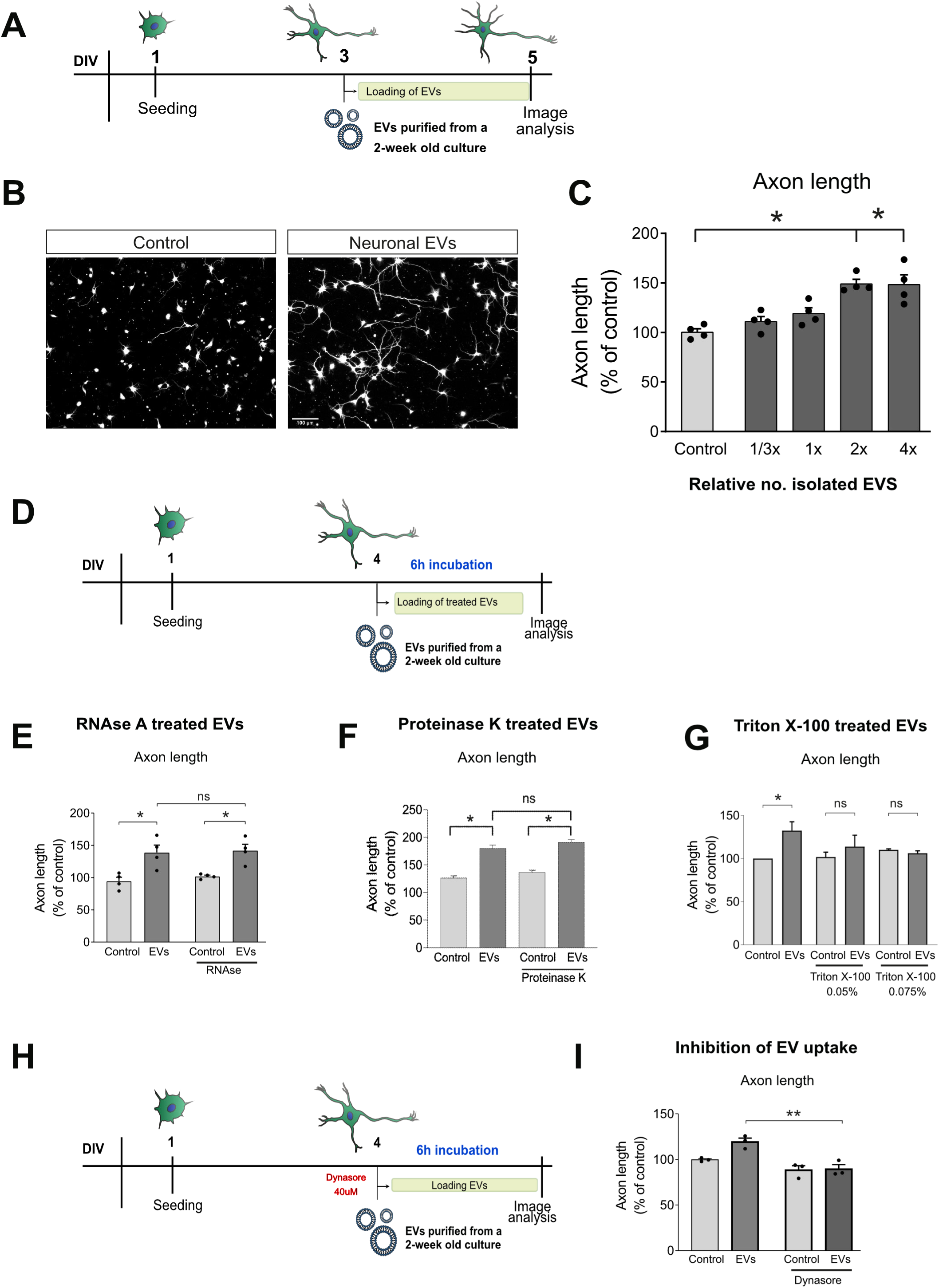
EVs derived from late-stage mouse primary cortical cultures influence the axon outgrowth of early-stage cortical neurons. **(A)** Diagrammatic representation of experimental model for the analysis of EV function in cortical development. **(B-C)** Addition of incremental number of EV particles isolated from late stage (2-week-old) cultures to early stage (DIV3) cortical neurons (48h incubation) led to an increase in axon length in a “dose-response” like manner, as compared to neurons treated with the EV elution buffer (DPBS) alone (acetylated tubulin, scale bar:100µm; mean±SEM; n=4) **(D)** Diagrammatic representation of experimental model for the analysis of EV cargo and its role in observed functional mechanisms. **(E)** Incubation of EV fraction with RNAse A (6h, following by addition of RNAse inhibitors) showed no effect on the growth-promoting actions of EVs on developing axons; n=4, while **(F)** incubation with proteinase K also failed to prevent the increase in axon growth from after EVs; n=3. **(G)** However, disruption of EV structure using Triton-X100 abolished the effect of EVs in axon growth of DIV4 neurons; n=3. **(H)** Diagrammatic representation of experimental model for the treatment with dynasore, an inhibitor of endocytic-associated protein dynamin to functionally disrupt EV transfer. **(I)** Neuronal primary cultures incubated with dynasore (40 µM) prevented the EV-mediated increase in axon growth (n=3). * p≤0.05 and ** p≤0.01.

To specifically link this developmental effect on axon growth to an EV-dependent process of inter-neuronal communication, it was important to systematically address several of the possible confounding factors that can be contributing to the observed effect. First, we investigated the potential role of RNA-containing structures that could be present in the extracellular space and may become associated with the EV surface. To test this, we devised an experimental model in which the isolated EV fraction was incubated with RNAse A for 30 min at 37° C to disrupt any potential ribonucleoproteins in the extra-vesicular space. First, we demonstrated the effectiveness of this experimental approach to degrade RNA (**Suppl Fig 2A**). Then, EVs previously incubated with RNAse A were added to cortical neurons at DIV4, and their development was evaluated 6 h later (**Fig 2D**). We found that RNAse addition did not prevent the EV-dependent increase in axon length (**Fig 2E**), confirming that extra-vesicular RNA in the extracellular domain is not mediating the effect in axon growth. A similar approach was employed to exclude potential contributions from extracellular or non-vesicular proteins associated with the EV fraction. Isolated EVs were treated with proteinase K for 30 min at 37 °C, a process capable of degrading protein-containing contaminants (**Suppl Fig 2B**), after which protease inhibitors were added to terminate the reaction. Importantly, EV fractions were still capable of promoting axonal growth after protease incubation (**Fig 2F**), demonstrating how intra-vesicular molecules mediate the increase in axonal growth. Finally, we incubated isolated EVs with increasing concentrations of X-100 Triton, aimed at disrupting both exosomes and microvesicles (61). Crucially, the disruption of EV integrity by detergent incubation removed the capacity for modulation of axonal growth (**Fig 2G**), without impacting overall neuronal integrity.

To further demonstrate active EV transfer in our experimental model we developed two independent approaches. First, we labelled isolated EVs with BioTracker^TM^ MemBright 560 Live Cell Dye and incubated these with developing cortical neurons as done in functional studies. As shown in **Suppl Fig 2C**, recipient cells acquired clear fluorescent staining, compared to dye-incubated mock-EV fractions (i.e. in absence of functional EVs). Finally, to functionally disrupt EV transfer, we incubated neuronal cultures with dynasore, an inhibitor of endocytic-associated protein dynamin (**Fig 2H**), which is an integral functional mediator in the uptake of EVs (68–70). As shown in **Fig 2I**, incubation of neuronal primary neurons with dynasore (40 μM) prevented the EV-mediated increase in axon growth. Collectively, these series of experiments allowed us to ascribe the observed functional effect to the EV-mediated inter-neuronal transfer of molecular content.

### Increase in axon growth via EVs is mediated by miR-99a

Following the demonstration that EV-dependent promotion of axonal growth was mediated by the transfer of intra-vesicular content, we focused on addressing the cellular and molecular processes behind this effect. Among potential mechanisms at play, a growing number of studies have identified non-coding RNAs as key components and mediators of EV function, particularly in cancer studies (71). In the CNS, their role has mainly focused on the effect of miRNAs in glia-neuron communication (72, 73).

Previous work in our lab has described the complex and highly specific sncRNA content of EVs released by cortical neurons, while simultaneously identifying several miRNAs with a conserved axonal localisation (31). By overlapping conserved axonal miRNAs reported in published datasets with the top 50 miRNAs detected in axons and EVs from primary cortical neurons, we identified a set of candidate miRNAs for further investigation in our EV studies (**Suppl Table 1**). As shown in **Fig 3A**, 13 miRNAs were identified from this strict and selective analytical workflow, including several with known axonal mechanisms, such as miR-9, miR-22, miR 26a, and/or reported as prognostic biomarkers in neurodegenerative processes, such as miR-181 (31, 33, 74, 75). From this analysis, we decided to focus on miR-99a, which is enriched in EVs but has not been previously shown as involved in developmental processes intrinsic to the axon.

**Figure 3.**
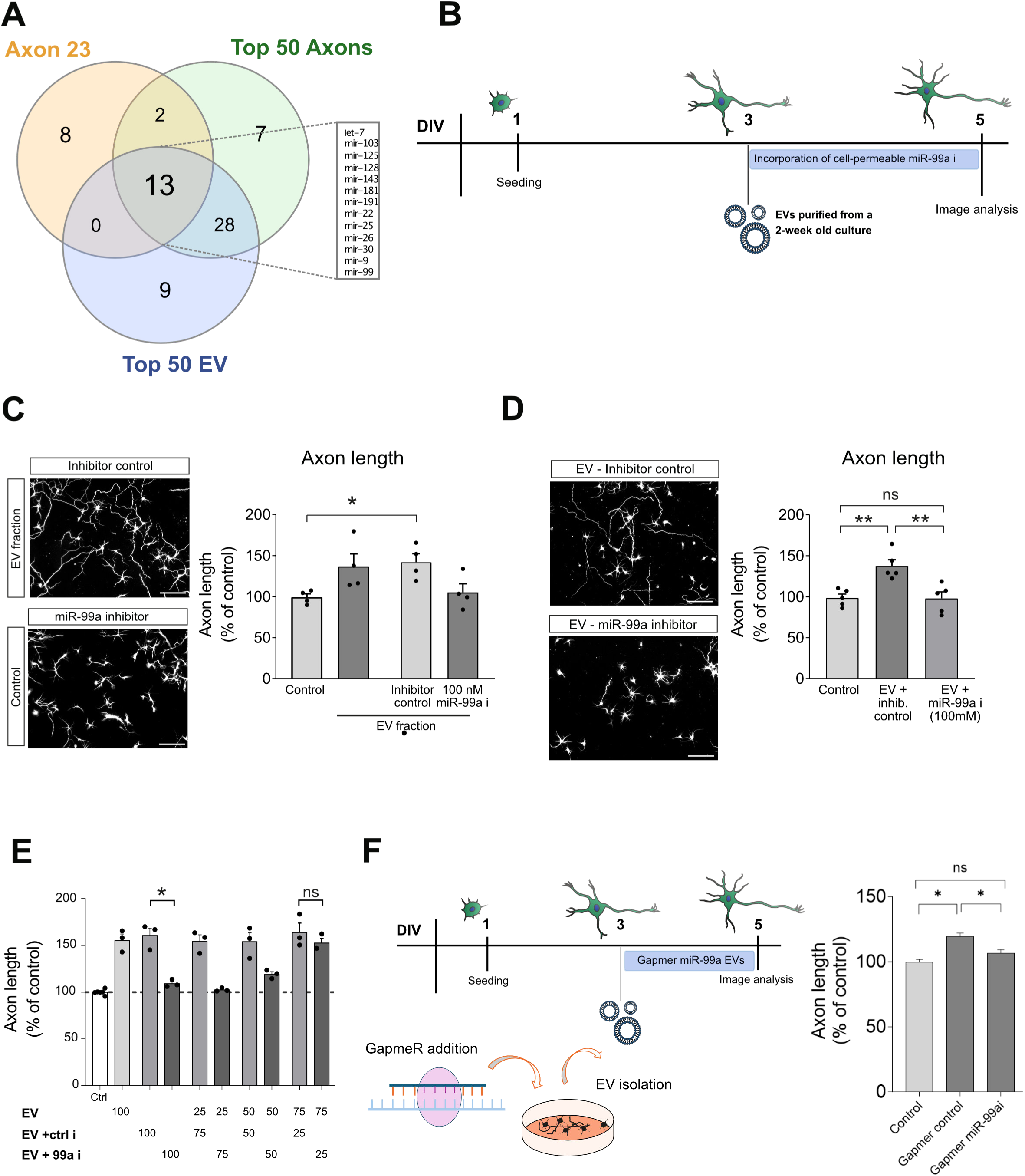
EV-dependent increase in axonal outgrowth is mediated by miR-99a. **(A)** Venn diagram depicting the subset of 13 miRNAs common to the top 50 most abundant in cortical neuron axon and EV samples, and the 23-miRNA signature in axons across models and species defined in Mesquita-Ribeiro et al. (2021). Following from this, miR-99a was selected for further investigation. **(B)** Diagrammatic representation of experimental model for the analysis of miR-99a function in EV-mediated axonal growth during neuron development. **(C)** Addition of a cell-permeable miR-99a inhibitor to neuronal cultures before incorporation of EVs prevents the EV-mediated increase in axonal growth. Whereas a cell-permeable non-targeting control showed no impact on EV growth promoting effects (n=4, scale 100 μm). **(D)** The incorporation of cell-permeable miR-99a inhibitor into EVs prior to addition to DIV 3 neuronal cultures, followed by SEC for removal of free oligonucleotide, also abolished the EV-dependent increase in axon growth as compared to the non-targeting control (n=5, scale 100 μm), suggesting the involvement of miR-99a cargo within EVs. **(E)** EV competition assay: a control population of cortical neuron-derived EVs was mixed with increasing amounts of separately isolated EVs containing the miR-99a inhibitor. Stepwise increases in the proportion of miR-99a inhibitor-containing EVs resulted in a progressive reduction in the axon growth-promoting effects observed with control EVs, supporting the role of miR-99a as a functional EV cargo in regulating axon growth (n=3). **(F)** Diagrammatic representation of experimental model for the analysis of axon length following incubation with EVs which had the maturation and processing of miR-99a blocked prior to EV isolation. EVs isolated after 48 hours of incubation with the pri-miR-99a GapmeR were applied to DIV3 primary neurons, these miR-99a-depleted EVs failed to promote axon growth, in contrast to EVs derived from cells treated with control GapmeR. * *p*≤0.05 and ** *p*≤0.01.

To test whether EV-derived miR-99a can modulate axon development in primary cortical neurons, we employed two experimental approaches. One was to incorporate a specific cell-permeable inhibitor of miR-99a to primary cortical neurons, prior to the addition of EVs, and to evaluate axon growth after 48 h, as done previously (**Fig 3B**). Inhibition of miR-99a in cortical neurons completely prevented the increase in axon length after addition of EVs (**Fig 3C**). In addition to this initial indication of a role for miR-99a, we devised a more compelling experimental approach, where the cell-permeable miR-99a inhibitor was incubated with the EVs before addition to the cortical culture, but only after performing a second EV isolation protocol to remove any remanent extra-vesicular inhibitor. Following this approach, the addition of EVs previously incubated with miR-99a inhibitor into primary cortical neurons also cancelled the EV-mediated increase in axon growth (**Fig 3D**). Importantly, in both experimental models the non-targeting miRNA inhibitor controls failed to prevent the EV-mediated increase in axonal growth.

Further evidence supporting the role of miR-99a as a functional EV cargo in regulating axon growth was obtained through EV competition assays. In these experiments, a control population of cortical neuron-derived EVs was mixed with increasing amounts of separately isolated EVs containing the miR-99a inhibitor. As shown in **Fig 3E**, stepwise increases in the proportion of miR-99a inhibitor-containing EVs resulted in a progressive reduction in the axon growth-promoting effects observed with control EVs.

To exclude the possibility that the effect of miR-99a on EV-mediated axon growth might involve EV-independent mechanisms, we assessed whether the miR-99a inhibitor could counteract the increase in axon length induced by protein-rich fractions eluting after the EV-enriched fractions during size exclusion chromatography. These fractions contain the majority of extracellular growth factors secreted by cortical neurons and were found to induce a modest but statistically significant increase in axon growth (**Suppl Fig 3A-B**). Notably, inhibition of miR-99a did not attenuate the axon growth-enhancing effect of the protein fraction (**Suppl Fig 3C-D**), reinforcing the EV-specific nature of miR-99a function in this context.

As an additional control for potential non-EV-related effects, we conducted experiments to rule out the possibility that contaminants present in cortical neuron culture media subjected to SEC were responsible for the observed outcomes. First, a “mock EV” SEC extraction was performed using fresh Neurobasal medium supplemented with B27, and qPCR analysis confirmed that miR-99a was undetectable in these fractions. Secondly, the isolated “mock EV” fractions did not induce any increase in axon growth when applied to cortical neurons, with or without B27 supplement, further supporting the specificity of the EV-mediated effects.

As a final approach to test the hypothesis that miR-99a functions as an EV cargo mediating axon growth changes in target neurons, we blocked the maturation and processing of miR-99a within cortical neurons prior to EV isolation. To achieve this, we designed an antisense LNA GapmeR (Qiagen) targeting pri-miR-99a in the nucleus, thereby preventing its maturation in the cytoplasm. Initial validation in the N2A neuron-like cell line demonstrated that EVs derived from GapmeR-treated cells exhibited a significant ∼ 20 % reduction in mature miR-99a content, as confirmed by qPCR. Applying this experimental strategy to primary cortical neurons, EVs were isolated after 48 hours of incubation with the pri-miR-99a GapmeR. When applied to DIV3 primary neurons, these miR-99a-depleted EVs failed to promote axon growth, in contrast to EVs derived from cells treated with control GapmeR constructs (**Fig 3F**). Collectively, the experiments presented in this section provide strong validation for the role of miR-99a as EV-cargo capable of regulating axon development.

### miR-99a regulates axon growth by targeting HS3ST2

The role of miR-99a in mediating the effects of EV on axonal length in culture points towards a cell mechanism for this miRNA in axonal growth and development. Indeed, neuron transfection with the miR-99a inhibitor in cortical neurons at 24 h after plating, produced a significant decrease in axonal growth evaluated 72 h later (**Suppl Fig 4A-B**). More importantly, with the aim to reproduce a model of increased delivery of this miRNA, transfection with a miR-99a mimic produced an increase in axon length compared to non-targeting control probes (**Suppl Fig 4C**). Collectively, this functional data provides direct confirmation for the role of miR-99a in the regulation of axonal outgrowth during development of neuron connectivity in cortical neuron cultures.

To investigate the molecular mechanism through which miR-99a regulates axon development, we searched for potential mRNA targets of miR-99a, using two extensively employed miRNA target prediction tools, TargetScan (76) and DIANA-microT-CDS (77) and the Human Protein Atlas to examine the reported expression of the target transcripts in ‘cerebral cortex’ (proteinatlas.org; (78). This concerted approach generated a list of 14 predicted targets that were shared between the two bioinformatic algorithms and detected in the brain (**Fig 4A and Suppl Table 1**). Among these top candidates, we focused on heparin sulfate-3-O-sulfotransferase 2 (Hs3st2) as a potentially relevant target of miR-99a in our model of axonal growth. This was based on previous reports showing expression of HS3ST2 in brain (56, 79) and cortical plates of mouse embryos (80). Furthermore, miR-99a binding site on *Hs3st2* 3’UTR is broadly conserved across vertebrate families and fully conserved in human (**Fig 4B**), further supporting the biological relevance of the miR-99a - *Hs3st2* interaction. Of note, this interaction has been already shown as a control to test the functionality of an adeno-associated virus-mediated microRNA delivery (81).

**Figure 4.**
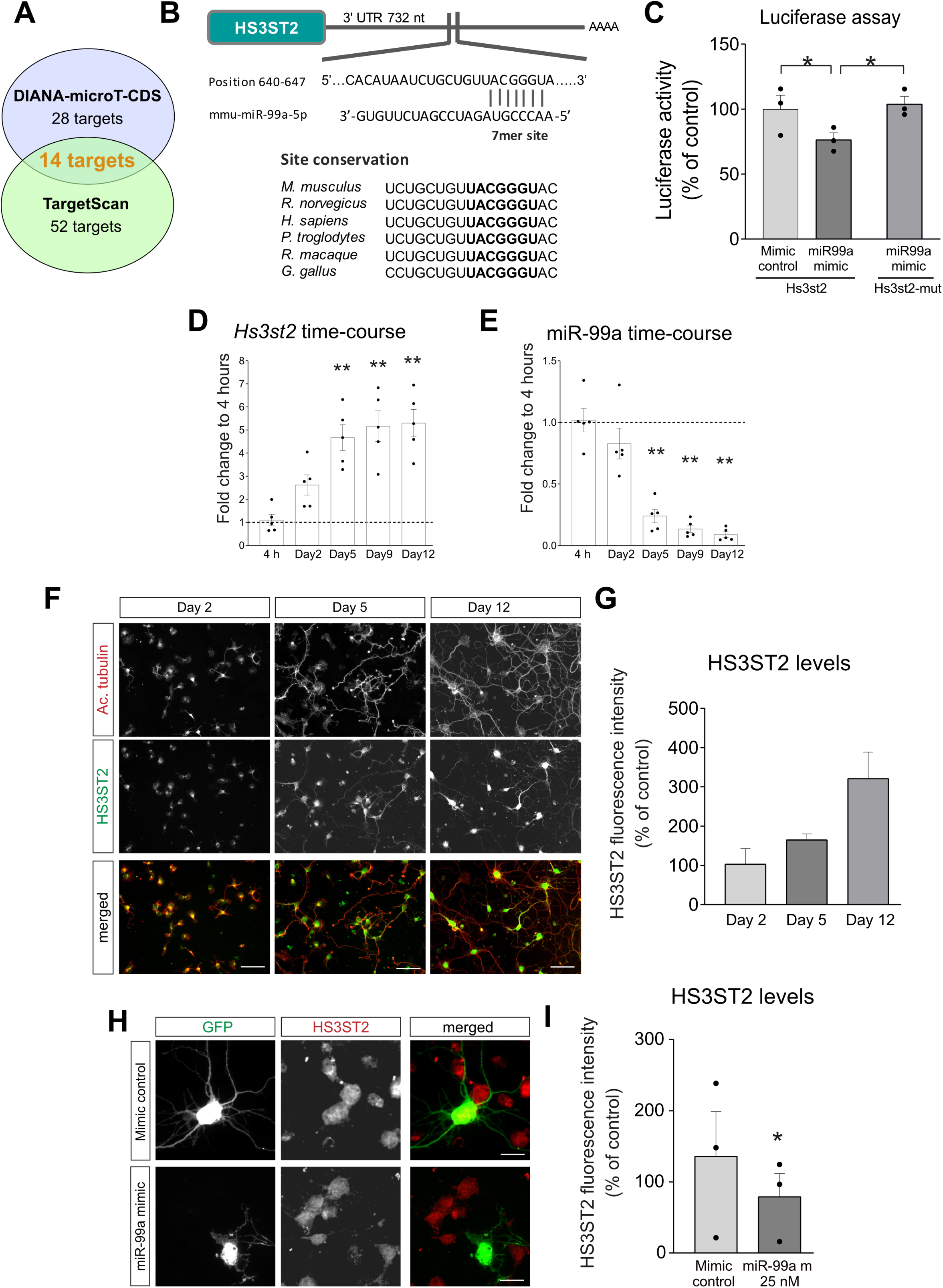
miR-99a targets heparan sulphate 3-O-sulfotransferase (*Hs3st2*) in mouse cortical primary neurons. **(A)** The bioinformatics approach for the identification of miR-99a targets used predictions from two independent algorithms – TargetScan and Diana Tools – to reach a subset of 14 common mRNAs illustrated. **(B)** Predicted miR-99a binding site (7mer) in the 3’UTR of *Hs3st2* transcript is broadly conserved in vertebrates, particularly across mammals. **(C)** Dual luciferase reporter assays where the miR-99a putative binding site within the *Hs3st2* 3’UTR was cloned into a firefly/renilla luciferase reporter plasmid. Results show a decrease in luciferase activity upon co-transfection with a miR-99a mimic in HEK-293T cells, which was abolished by mutation of the binding site, thus providing evidence for miR-99a – *Hs3st2* interaction (n=3). RT-qPCR of *Hs3st2* **(D)** and miR-99a levels **(E)** in cortical neurons at 4h, DIV2, DIV5, DIV9 and DIV12 show opposite expression profiles over development in culture, suggesting a miR-mRNA target co-regulation (ddCt method, fold change to 4h; mean±SEM, n=5). **(F-G)** HS3ST2 protein levels replicate the expression profile of its transcript, as measured by fluorescence intensity of the protein by immunofluorescence of cortical neurons (HS3ST3: green, acetylated tubulin: red) at DIV2, 5 and 12 (expressed as percentage of DIV2, mean±SEM; n=3, scale 100 μm). **(H-I)** Transfection of DIV2 neurons with a miR-99a mimic leads to a decrease of HS3ST2 protein levels detected by immunofluorescence, 72h after transfection, indicating that miR-99a can directly regulate HS3ST2 in developing cortical neurons (data presented as percentage of control, mean±SEM; n=3, scale 25 μm). * *p*≤0.05 and ** *p*≤0.01.

To assess whether miR-99a regulates *Hs3st2* expression, we first generated a dual luciferase reporter construct by inserting the predicted miR-99a binding site from the *Hs3st2* 3′ UTR downstream of the firefly luciferase gene, co-expressed with a constitutively active Renilla luciferase gene for normalisation. The reporter plasmid was co-transfected into HEK293T cells alongside either a miR-99a mimic or a non-targeting control mimic, and luciferase activity was subsequently measured. The presence of the miR-99a mimic led to significant repression of the reporter, with a 23.6% decrease in the firefly/Renilla activity ratio compared to the control (**Fig 4C**). Importantly, site-directed mutagenesis of four nucleotides within the seed region of the predicted miR-99a binding site abrogated this repression, thereby confirming the specificity of miR-99a targeting to the *Hs3st2* 3′ UTR (**Fig 4C**).

To further support the role of HS3ST2 in our model, we examined its expression in primary mouse cortical neuron cultures using qPCR. As shown in **Fig 4D**, *Hs3st2* transcripts were detectable as early as 4 hours post-plating, with expression gradually increasing and stabilising through to 12 days in vitro (DIV12). In contrast, miR-99a expression showed an inverse pattern (**Fig 4E**), progressively decreasing with time in culture, consistent with a regulatory interaction between the miRNA and its target. At the protein level, HS3ST2 expression was evaluated by quantitative immunostaining and normalised to DIV2 levels, revealing a parallel temporal profile to its transcript. By DIV12, HS3ST2 protein levels had increased approximately 7.5-fold relative to DIV2 (**Fig 4F-G**). Additional validation was obtained by modulating miR-99a levels in primary neurons: overexpression of the miR-99a mimic resulted in reduced HS3ST2 protein levels (**Fig 4H-I**), mimicking the functional effect mediated by mir-99a delivery via EV-cargo.

Having established the ability of miR-99a to regulate *Hs3st2* expression, we next investigated the previously uncharacterised role of HS3ST2 in axon development. To this end, we performed both loss- and gain-of-function experiments (**Fig 5A**). Loss-of-function was achieved via transient transfection of primary cortical neurons at DIV2 with either *Hs3st2*-targeting siRNA (50 nM) or a non-targeting control. Gain-of-function was carried out by transfection with a construct overexpressing HS3ST2. Overexpression of HS3ST2 led to a significant reduction in axon length compared to the empty vector control (**Fig 5B**). Conversely, silencing of *Hs3st2* resulted in axons that were, on average, 39.0% longer than those in control neurons (**Fig 5C**).

**Figure 5.**
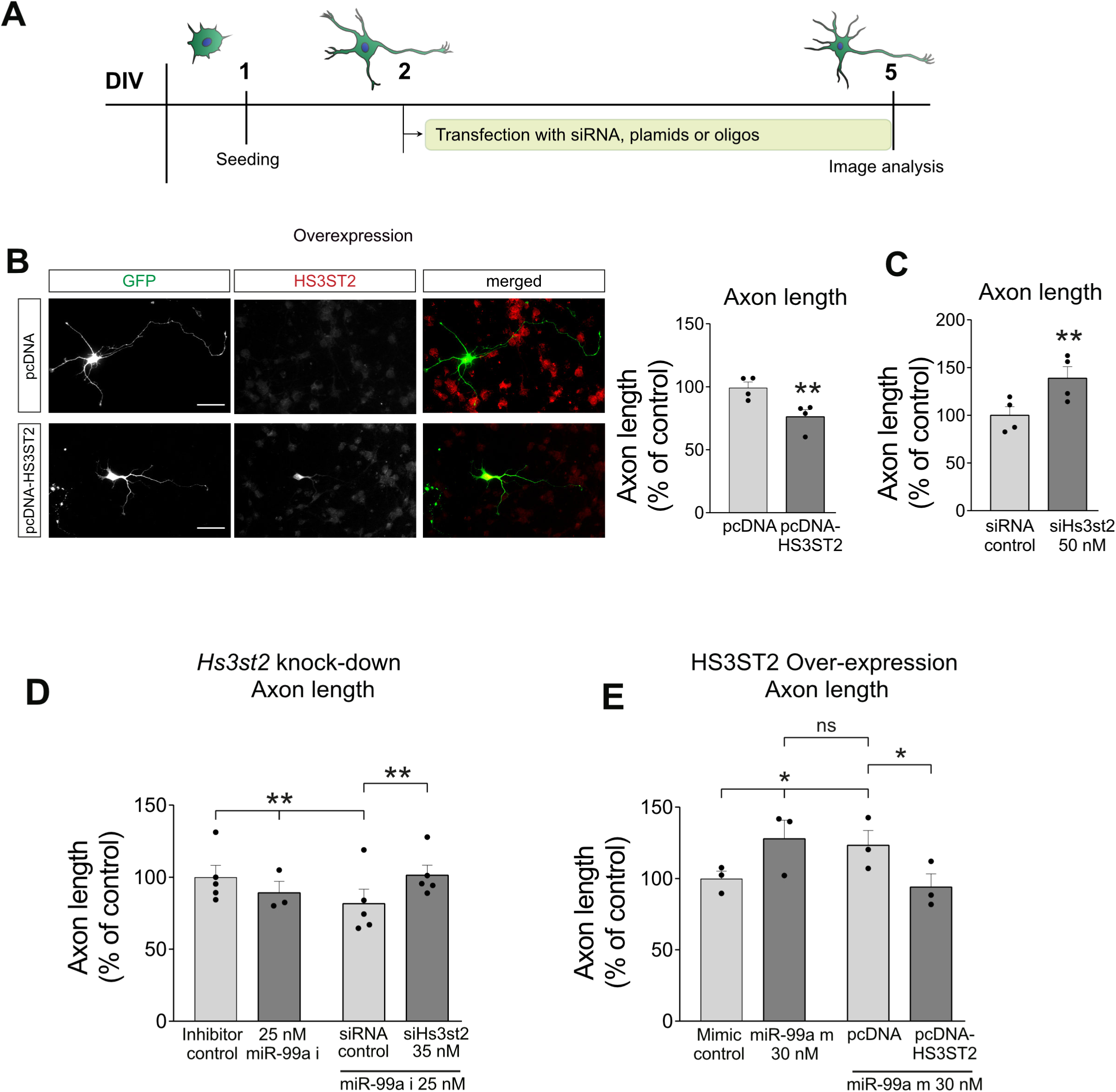
miR-99a promotes outgrowth in cortical axons via regulation of HS3ST2. Top panel showing a diagrammatic representation of experimental model for the addition of siRNA, plasmids or oligos as required. **(A)** Co-transfection of pGFP (as transfection reporter) and pcDNA-HS3ST2 resulted in a decrease in the length of axons as compared to the empty vector (n=4, scale 40 μm), **(B)** whereas a significant increase in axon length was observed upon transfection of DIV2 neurons with siHs3st2 (co-transfected with pGFP); n=4. **(C-D)** DIV2 neurons were co-transfected with miR-99a inhibitors or mimics and siHs3st2 or pc-HS3ST2, respectively, and their axons measured 72h later. Hs3st2 knockdown **(C)** or overexpression **(D)** rescued the functional effects in axonal outgrowth of miR-99a inhibitors or mimics, respectively, taking axon length measurements back to control levels (GFP:green, HS3ST2: red; data presented as percentage of control, mean±SEM; n=5 and 3 respectively). * *p*≤0.05 and ** *p*≤0.01.

To further explore the functional link between miR-99a and HS3ST2 in axon growth, we conducted rescue experiments. Knockdown of HS3ST2 reversed the decrease in axon length typically induced by miR-99a inhibition (**Fig 5D**), while overexpression of HS3ST2 (pcDNA construct lacking the 3’ UTR) counteracted the axon growth-promoting effect of the miR-99a mimic, restoring axon length to control levels (**Fig 5E**). The rationale behind these experiments was that modulating HS3ST2 expression should offset the phenotypic effects of altering miR-99a levels, thereby functionally linking the two. It is important to note, however, that miRNA-mRNA “competition/rescue” paradigms do not operate in a simple binary, receptor-ligand manner. Rather, they reflect more nuanced, gradient-like effects on downstream biological outcomes, such as axon growth. Despite this complexity, our findings provide strong evidence for a novel mechanism of axonal growth regulation mediated by miR-99a-dependent targeting of HS3ST2.

### EVs regulate axonal growth via miR-99a targeting of *Hs3st2*

Having demonstrated that miR-99a regulates axonal growth via targeting of *Hs3st2*, we next investigated whether this interaction underlies the axon growth effects observed with EV treatment. To address this, we implemented a “double-rescue” experimental strategy to test whether knockdown of *Hs3st2* could offset the inhibitory effects of miR-99a suppression, and the consequent increase in HS3ST2 protein levels, on EV-mediated axon elongation. As in previous experiments, transfections were performed 24 hours after plating, and axon length was assessed 72 hours later. Consistent with prior results, the axon growth-promoting effect of EVs was abolished by specific inhibition of miR-99a (**Fig 6B**), but not by a non-targeting inhibitor control. Importantly, co-transfection with *Hs3st2*-targeting siRNA restored the EV-induced increase in axon length, supporting the notion that miR-99a facilitates axon growth through repression of HS3ST2 in recipient neurons.

**Figure 6.**
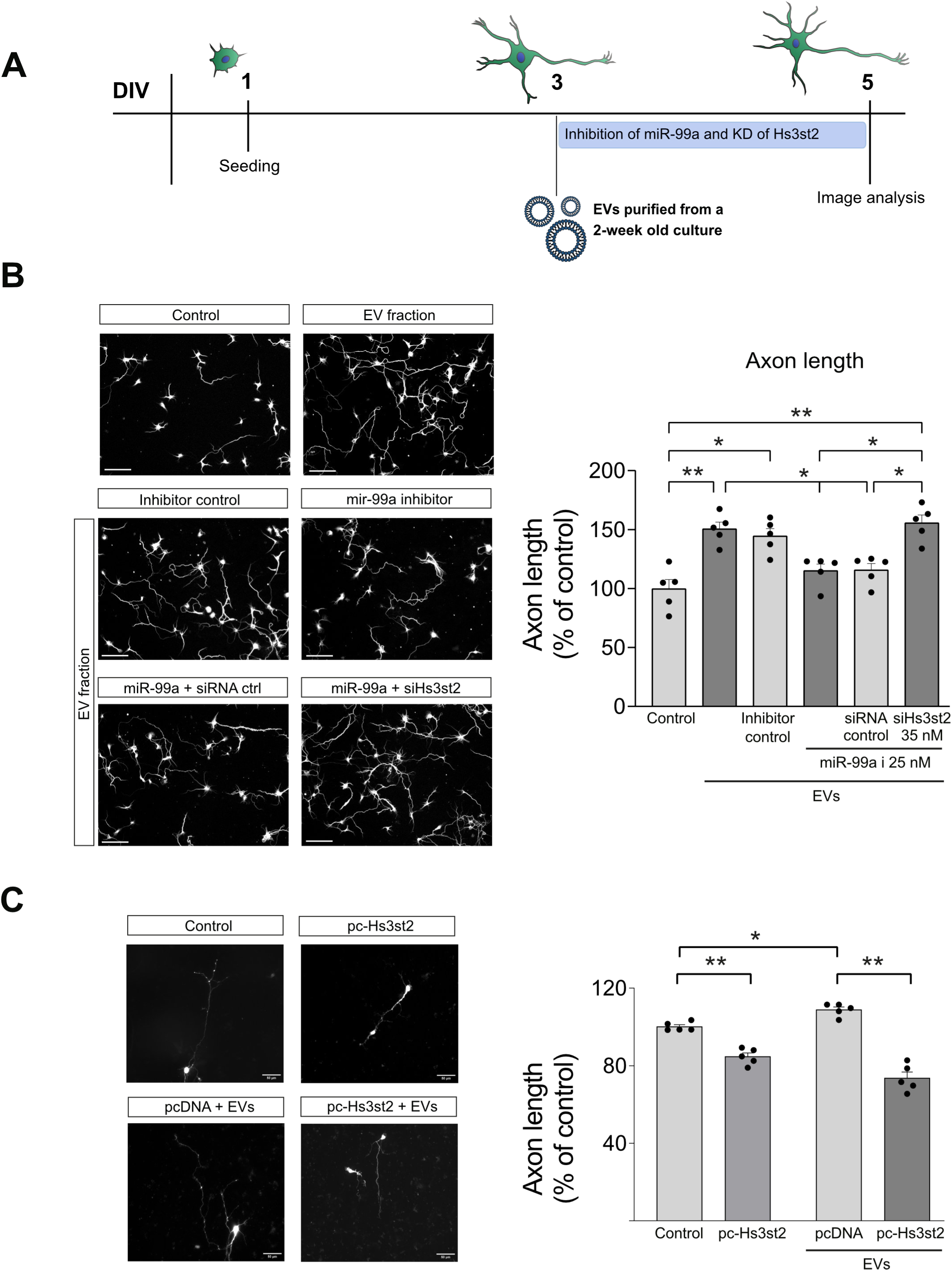
EV-derived miR-99a controls axonal development in cortical neurons, in a mechanism involving the regulation of HS3ST2. **(A)** Diagrammatic representation of the functional rescue of axon length upon Hs3st2 knockdown in neurons at DIV3 and following addition of EVs where miR-99a has been inhibited. **(B)** In neurons incubated with EVs plus cell-permeable miR-99a inhibitor, knocking-down HS3ST2 protein using cell-permeable siHs3st2 abolished the previously observed miR-99a rescue of EV-mediated increase in axon growth, restoring axon length to the levels measured in neurons treated with EVs only (acetylated tubulin, scale 100 μM ; n=5, data presented as percentage of control, mean±SEM). **(C)** In neurons overexpressing a miR-99a-resistant *HS3ST2* construct (pc-HS3ST2), which lacks the 3′ UTR, EV treatment continued to promote axon elongation in the empty plasmid control pcDNA, but this effect was abolished when neurons overexpressed pc-HS3ST2, demonstrating that repression of HS3ST2 by EV-delivered miR-99a is essential for mediating its pro-axon growth activity (n=5). * *p*≤0.05 and ** *p*≤0.01.

To further validate that EV-derived miR-99a exerts its effects via direct targeting of *Hs3st2* in recipient cells, we overexpressed the *HS3ST2* construct (pcDNA-HS3ST2), lacking the 3′ UTR. As shown in **Fig 6C**, EV treatment continued to promote axon elongation even in the high-density cultures that are required for neuronal transfection. This effect was abolished when neurons overexpressed the miR-99a-resistant *HS3ST2* construct, demonstrating that repression of HS3ST2 by EV-delivered miR-99a is essential for mediating its pro-axon growth activity.

### EVs control of axonal growth is local to the axon compartment

Our findings demonstrate that neuron-derived EVs influence axon development; however, standard culture systems cannot define where this effect occurs. To determine if EV-mediated mechanisms are local to specific sub-cellular domains within the complex morphological structure of a neuron, we employed compartmentalised microfluidic chambers, which allow the fluidic isolation of axons from cell bodies. In this experimental model, neurons are seeded on one side of the device and axons allowed to grow onto the opposite side. Addition of EVs to the axon side of the chambers on day 6 of culture significantly promoted axonal growth (**Fig 7B**), as shown previously in normal culture dishes. However, this growth-promoting effect on the axonal side was not seen when the EVs were added exclusively to the soma side of the microfluidic device (**Fig 7C**). Overall, the use of this model points towards a cellular mechanism where EVs can target axonal growth by regulating local translation processes locally. To confirm this, we analysed HS3ST2 protein levels in different subcellular domains after specific addition of EVs to either the somatic or axonal compartments in microfluidic chambers. As shown in **Fig 7D**, EVs added to the axonal compartment significantly reduced HS3ST2 protein levels in growth cones, consistent with results from conventional cultures, but had no effect in cell bodies. When EVs were applied to the somatic compartment instead, HS3ST2 levels in growth cones remained unchanged (**Fig 7E**). As with the functional axon growth data, these results point towards a direct effect of EVs on the axonal subdomain, locally controlling cellular function via targeting HS3ST2 expression.

**Figure 7.**
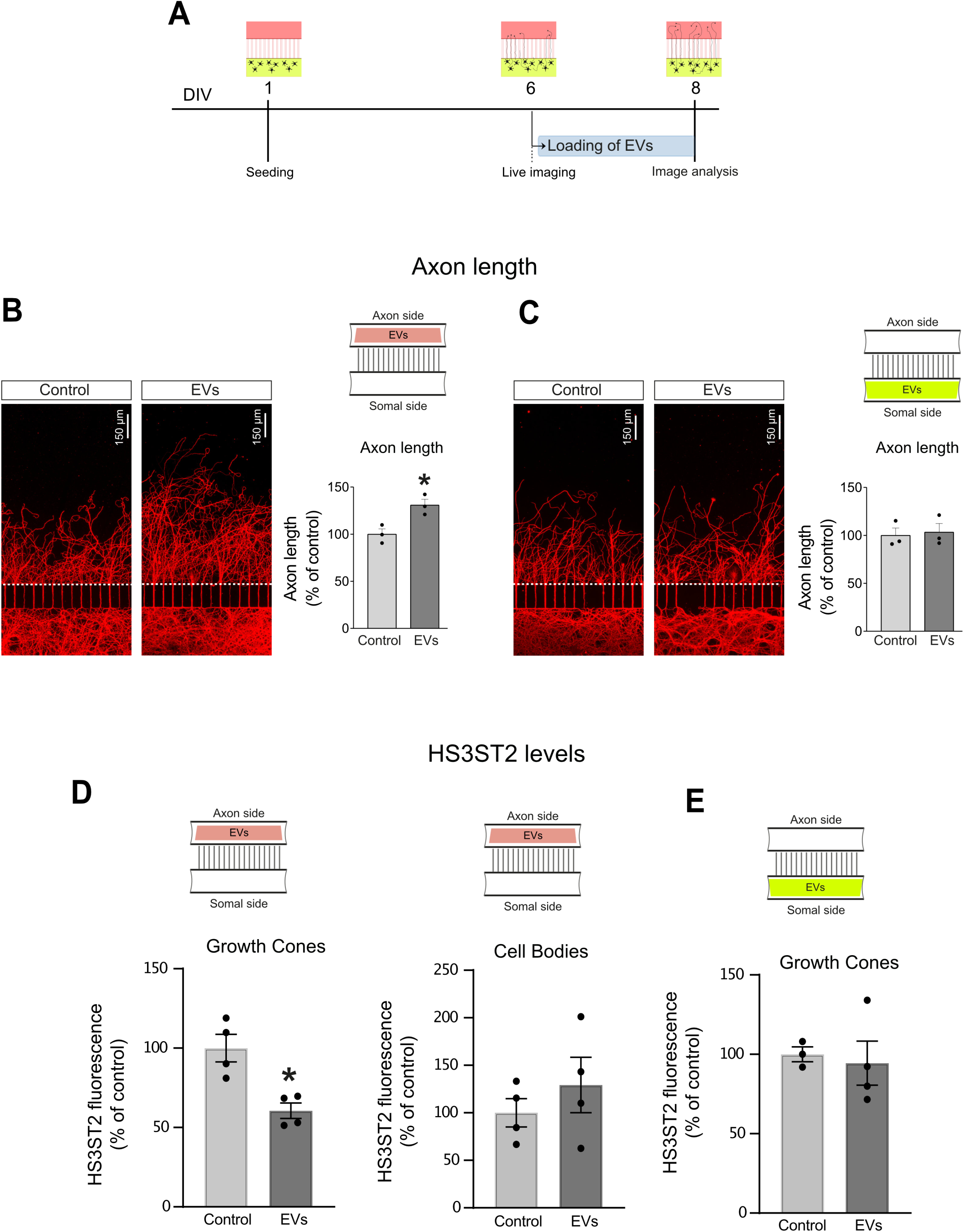
Neuron-derived EVs act locally in the axon to promote axonal outgrowth. **(A)** Using a compartmentalised microfluidic culture model in which axons of cortical neurons are fluidically isolated from the somatodendritic compartment, **(B)** EVs were added to the axonal side of the microfluidic chamber after axons have extended into the axonal compartment (DIV6), leading to an increase in axon length compared to controls (DPBS, 48h incubation, n=3). **(C)** The EV-mediated increase in axonal length was not observed when EVs were added to the somatodendritic compartment, suggesting a local growth promoting effect of neuron-derived EVs in developing axons (n=3). **(D)** Loading of EVs onto the axonal compartment alone resulted in a significant reduction of HS3ST2 expression levels in the axonal growth cones as quantified by immunofluorescence, but not in the neuronal somas (n=3). **(E)** When EVs were added into the somatodendritic compartment alone, no significant changes in HS3ST2 expression were observed in the growth cones (n=3), (acetylated tubulin:red; scale bar:150μm; data presented as percentage of control, mean±SEM). * *p*≤0.05.

## DISCUSSION

The findings presented in this study highlight the critical role of EV-mediated inter-neuronal communication in regulating local axonal translation through the delivery of microRNAs. Our data demonstrate that subcellular modulation of gene expression can exert rapid and spatially restricted effects on axonal dynamics, thereby influencing key aspects of neurodevelopment. Importantly, this work underscores the potential of EVs to orchestrate axon growth by modulating the finely tuned spatial and temporal expression of *HS3ST* family genes, molecular regulators increasingly recognised for their roles in both developmental processes and neurodegenerative conditions.

### Inter-neuronal communication via EVs

Our findings contribute to the growing understanding that EV biogenesis can be dynamically regulated by different cell types according to their physiological state, resulting in the release of vesicles with distinct lipid, protein, and nucleic acid profiles (12). This EV-mediated intercellular communication, operating through both paracrine and autocrine mechanisms, relies on vesicle docking and internalisation, either via endocytosis or direct membrane fusion with recipient cells. However, the downstream cellular pathways triggered by EV uptake remain incompletely characterised, and the specific functional consequences of cargo delivery, particularly in the context of neural development, are only beginning to be elucidated.

In the nervous system, early work on cell-cell EV transfer focused on their role as mediators of <u>glia-neuron</u> communication (82–84), in a process that was shown to impact both neuronal network degeneration (72, 85) and neuronal plasticity (73). More recently, EV delivery of miR-126a by muscle cells was demonstrated to control TDP-43 local translation in axons, regulating neuromuscular junction maintenance (Ionescu et al., 2025). In <u>neuron-to-neuron</u> interactions, presynaptic exosome release was shown to regulate postsynaptic retrograde signalling (19), while the trans-synaptic transfer of EVs was revealed to specifically target recipient neurons (20), regulating spontaneous activity and spine formation (50, 51). Moreover, the inter-neuronal exchange of synaptobrevin via EVs was demonstrated to rapidly integrate into synaptic vesicle cycle of host neurons and selectively augment inhibitory neurotransmission (23). Neuron derived EVs can also promote neural differentiation of adipose-derived stem cells, in a model that could be applied in the prevention of peripheral nerve injury degeneration (64). Interestingly, a recent study showed how neuron-derived EVs are enriched in proteins linked to “nervous system development networks” and their injection into the lateral ventricle of postnatal mice could increase neural cell proliferation (22). Although early EV studies also demonstrated the release of exosomes by cultured cortical neurons (17, 18), and in the case of neuroblastoma models how activity-associated miRNAs could be released in exosomes after depolarization (21), evidence of a role for EV-RNA cargo in functional neuronal outputs has been limited to a recent study on the BDNF-dependent control of dendritogenesis and synapse maturation (86). Our study provides some of the earliest evidence for EV-mediated neuron-to-neuron transfer of a specific miRNA capable of regulating local gene expression to influence axon development in central nervous system neurons. The targeted delivery of miRNAs to axonal subcellular compartments via EVs represents a powerful mechanism for modulating neuronal function, particularly given the typically low copy number of miRNAs within individual EVs. Based on our experimental model, it is possible to conceive a mechanism in which the developmentally regulated decline in endogenous miR-99a levels (as shown by whole-cell qPCR in **Fig 4E**) may be locally counterbalanced by EV-mediated miR-99a delivery to axons. This, in turn, could suppress the translation of increasingly abundant *Hs3st2* mRNA (also shown in **Fig 4E**), thereby promoting axon elongation. In effect, we demonstrate how sub-cellular control of local translation has the potential to impact cellular “neighbourhoods” of axon growth cones independent of the regulatory environment in the host’s cell body domain. Our demonstration of subcellular miRNA function contributes to reconciling the apparent discrepancy between stoichiometric analyses of EV miRNA content and their predicted functional relevance. Notably, early work by Chevillet et al. (87) estimated that, on average, exosomes contain far less than one copy of a given miRNA, leading to the prediction that most individual vesicles may not carry biologically meaningful miRNA quantities. However, single-molecule studies of endogenous *β-actin* mRNA in axons have shown that even functionally relevant transcripts are present at very low copy numbers, fewer than four per axon growth cone in most cases (88). Together, these analyses underline the axon growth cone as a functionally meaningful cellular vessel for EV regulation of gene expression. Moreover, extensive evidence has shown that thousands of mRNA species undergo local translation within axonal and synaptic regions (89–92), underscoring the significant contribution of local protein synthesis to the proteomic landscape of these compartments. In this context, our study identifies a novel mechanism for subcellular protein regulation via miRNA activity that operates independently of somatic control, highlighting the nuanced and spatially restricted nature of gene expression in developing neurons.

### miR-99a in cell regulatory mechanisms

The miR-99 family comprises miR-99a, miR-99b, and miR-100, which share a conserved 7-nucleotide seed region. Notably, miR-99a and miR-99b differ by only a single nucleotide in their mature sequences, while miR-100 differs by four nucleotides. Although the expression of these miRNAs is tightly regulated, numerous studies have implicated them in the progression of nearly all major human cancers, functioning either as oncogenic miRNAs (oncomiRs) or tumour suppressors (93). This duality likely reflects their capacity to target a broad array of transcripts involved in cell proliferation, differentiation, and survival. In addition, their role in tumour biology is further reinforced by evidence that they influence immune cell activity and differentiation, thereby modulating the tumour microenvironment and facilitating cancer progression (93).

In the CNS, the genomic location of miR-99a on chromosome 21 contributes to its overexpression, by at least 50%, in the foetal hippocampus of individuals with Down syndrome compared to controls (94), while recent studies have shown how overexpression of miR99a after intra-hippocampal lentiviral delivery resulted in reversal learning impairment in mice, without impacting motor function or anxiety (41). These findings suggest that miR-99a can exert a significant influence on neuronal development and cognitive function by modulating the expression of multiple genes implicated in synaptic plasticity, neuronal differentiation, survival, and immune responses (41).

Within neuronal mechanisms, despite the identification of miR-99a in a short list of conserved axonal miRNAs (31), its function in neuron development and axonal mechanisms had not been previously explored. Interestingly, miR-99a, together with miR-29a and miR-125a, have been shown to be released by synaptosomes under physiological stimulation and were proposed as part of miRNA secretion processes that could modulate nerve cell terminals (45). Recent work has shown that EV-delivered miR-100, a member of the miR-99 family, promotes endometrial stromal cell proliferation, invasion, and migration through the inhibition of *HS3ST2* (95). This highlights a striking conservation of molecular mechanisms that govern cell growth and invasiveness, features also relevant to axonal extension and navigation. In our study, we identify a novel role for miR-99a in neuronal development, demonstrating that its expression and EV-mediated transfer can modulate axon growth via the specific targeting of *HS3ST2*. These findings offer new insight into the role of miR-99a as a regulator of inter-neuronal communication and highlight *HS3ST2* as an emerging key player in cell growth regulation, neuronal function, and potentially, neurodegenerative processes.

### Heparan sulfates and HS3ST2

Structurally organised as repetitive linear disaccharide sequences, heparan sulfates (HS) belong to the family of sulphated glycosaminoglycans (GAGs) and are the glycanic constituents of HS proteoglycans (HSPGs). Within their complex biosynthetic machinery, sulfotransferases integrate sulfate groups into the elongating sugar chain, providing high structural complexity to HS and the capacity to form differently sulphated clusters among each chain. As a result, the specific regulation of the HS metabolic machinery in each cell type and subcellular domain can produce different sulfate signatures, which can affect the strength of HS cellular interactions with a large number of proteins or peptides (generically known as heparin binding proteins and including growth factors and cytokines among many).

The HS3ST2 enzyme is mainly expressed in neurons, where it generates rare 3-O-sulphated domains in HS. Although 3-O sulfation constitutes a relatively rare modification, the HS3STs are one of the largest families involved in HS biosynthesis, which suggests an important role in cellular function. Given their various HS chains and localisation (cell surface or ECM) HSPGs have been proposed to regulate signalling processes in multiple ways, including cell autonomously as receptors or co-receptors, as recruiters to lipid rafts, by regulating receptor membrane trafficking, or by controlling ligand secretion, while non-cell autonomous mechanisms also include the control of distribution and composition of the ECM (56). Overall, the large variability of HS chains that can be generated led to the proposal of the existence of a “sugar code”, where tissue-specific HS modifications could orchestrate cellular signalling programmes during nervous system development, including neurogenesis and axon pathfinding/development (56, 79, 96, 97). Interestingly, the role of HS3ST2 has grown beyond that seen during development, with increased levels found in the brains of AD patients and a proposed role in the molecular mechanisms leading to the abnormal phosphorylation and aggregation of tau (57, 98) and neurodegeneration (99).

Despite the recognised importance that regulatory mechanisms controlling modifications of HSPGs have gained so far, and the critical role that has been ascribed to 3-O sulfation (79), the cellular mechanisms controlling the complex spatial and temporal expression of HS3STs genes is largely unknown. Our findings have uncovered a novel role for HS3ST2 in the localised control of neuron development, and crucially, provided a molecular mechanism that regulates its localised expression in subcellular domains within the axon. The importance of this is manifold, providing evidence for the capacity for localised control of neuronal interactions within a changing ECM environment, and via the autocrine control of gene expression through EV-mediated delivery of miRNAs.

### Conclusions

Our findings demonstrate that EVs can modulate axon development through a miRNA-dependent mechanism, enabling the fine-tuning of local protein synthesis in a manner that operates independently of somatic control. This work adds to the growing body of evidence supporting the role of miRNAs in shaping subcellular protein composition, thereby influencing coordinated cellular behaviours during neural development. Future studies should explore how neurons dynamically regulate EV miRNA cargo across different developmental stages and in response to environmental cues, as well as investigate how this mode of intercellular communication contributes to the establishment and refinement of neuronal connectivity *in vivo*.

## Aknowledgements

The authors thank the School of Life Sciences Imaging and Microscopy (SLIM) Facility and the Nanoscale and Microscale Research Centre (nmRC) at the University of Nottingham for instrumentation access, and Denise McLean for technical assistance with sample preparation.

## Figure Legends

**Supplementary Figure 1.**
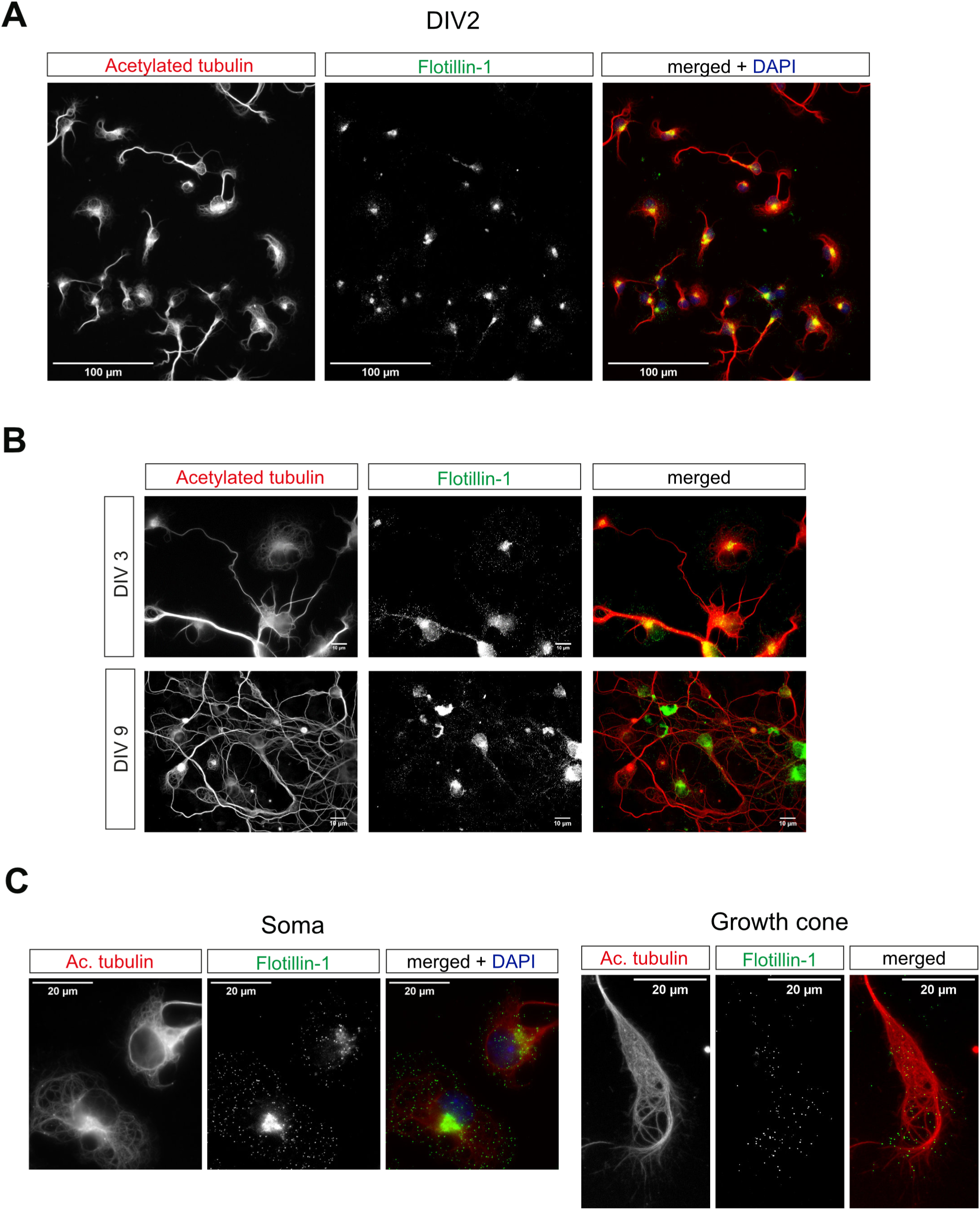
Mouse primary cortical neurons generate extracellular vesicles (EVs) in culture. **(A)** Immunostaining of primary cortical neurons with flotilin-1 (green; scale bar:100μm) demonstrates an **(B)** increase in expression over development (DIV3 vs DIV9; scale bar 100μm) and distribution in both **(C)** the cell soma and axon/growth cone structures (acetylated tubulin, red; DAPI, blue; scale bar:20μm).

**Supplementary Figure 2.**
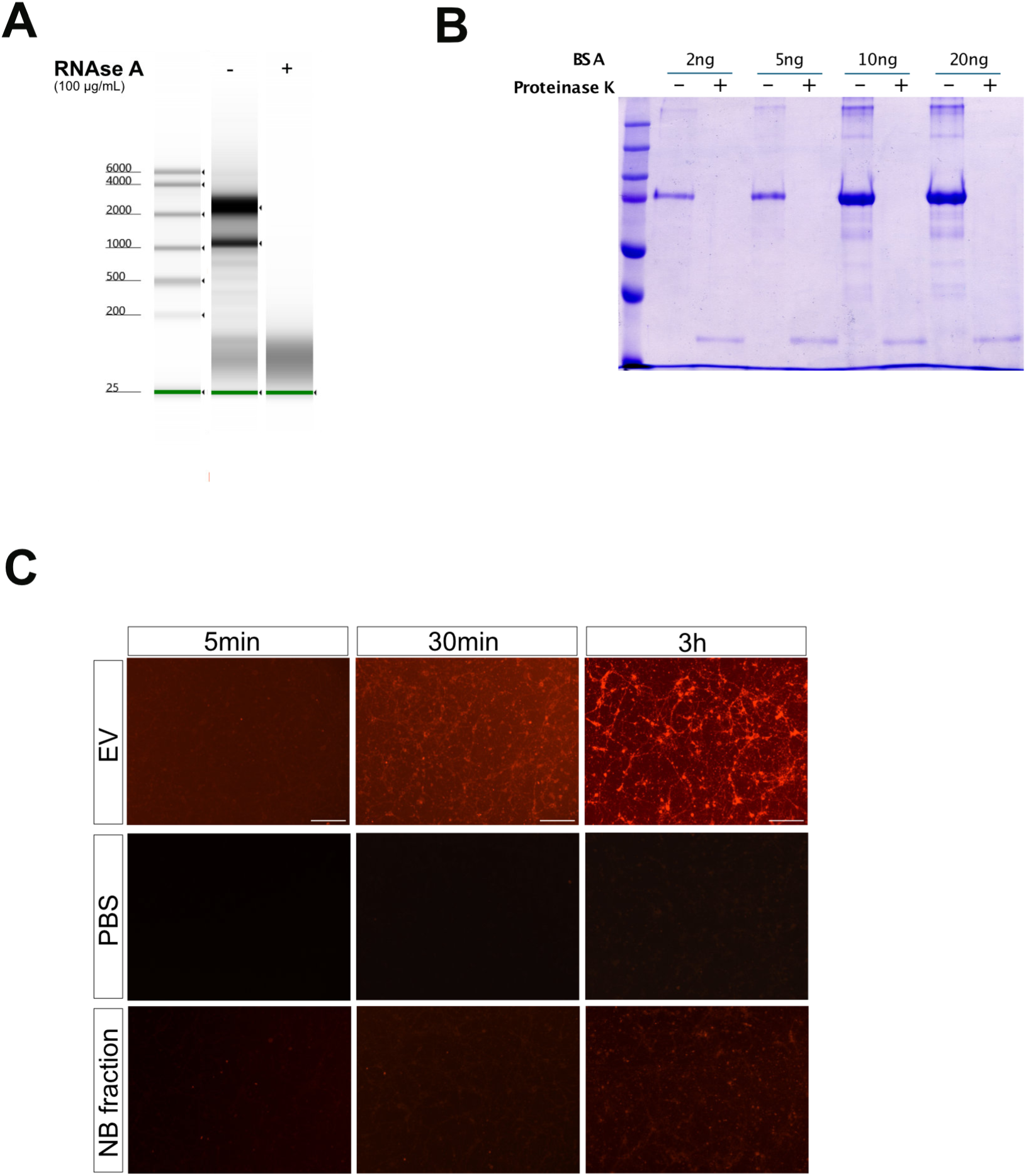
Validation of enzymatic degradation and EV transfer assays used in this study. **(A)** RNA incubated with 100µg/mL RNAse A for 30 min at 37 °C. Untreated and RNase A treated samples were analysed using the High Sensitivity RNA ScreenTape, using the Agilent 4200 TapeStation System. Results demonstrate the effectiveness of this experimental approach to degrade RNA as shown by the loss of the 28 and 18s RNA fragments. **(B)** Various bovine serum albumin (BSA) concentrations (2-20 ng) treated with 50 μg/mL proteinase K for 30 min at 37 °C, ran by SDS-PAGE and stained by Coomassie staining to visualise protein degradation. Results show that the proteinase K successfully degrades BSA protein. **(C)** To demonstrate active EV transfer in our experimental model isolated EVs were labelled with 200 nM BioTracker^TM^ MemBright 560 Live Cell Dye, isolated using size exclusion chromatography and incubated with developing cortical neurons. Recipient cells acquired clear fluorescent staining which increased over time (labelled EV), compared to the controls; PBS alone or mock EV fractions (NB Fraction) from fresh media that were processed similarly to EVs.

**Supplementary Figure 3.**
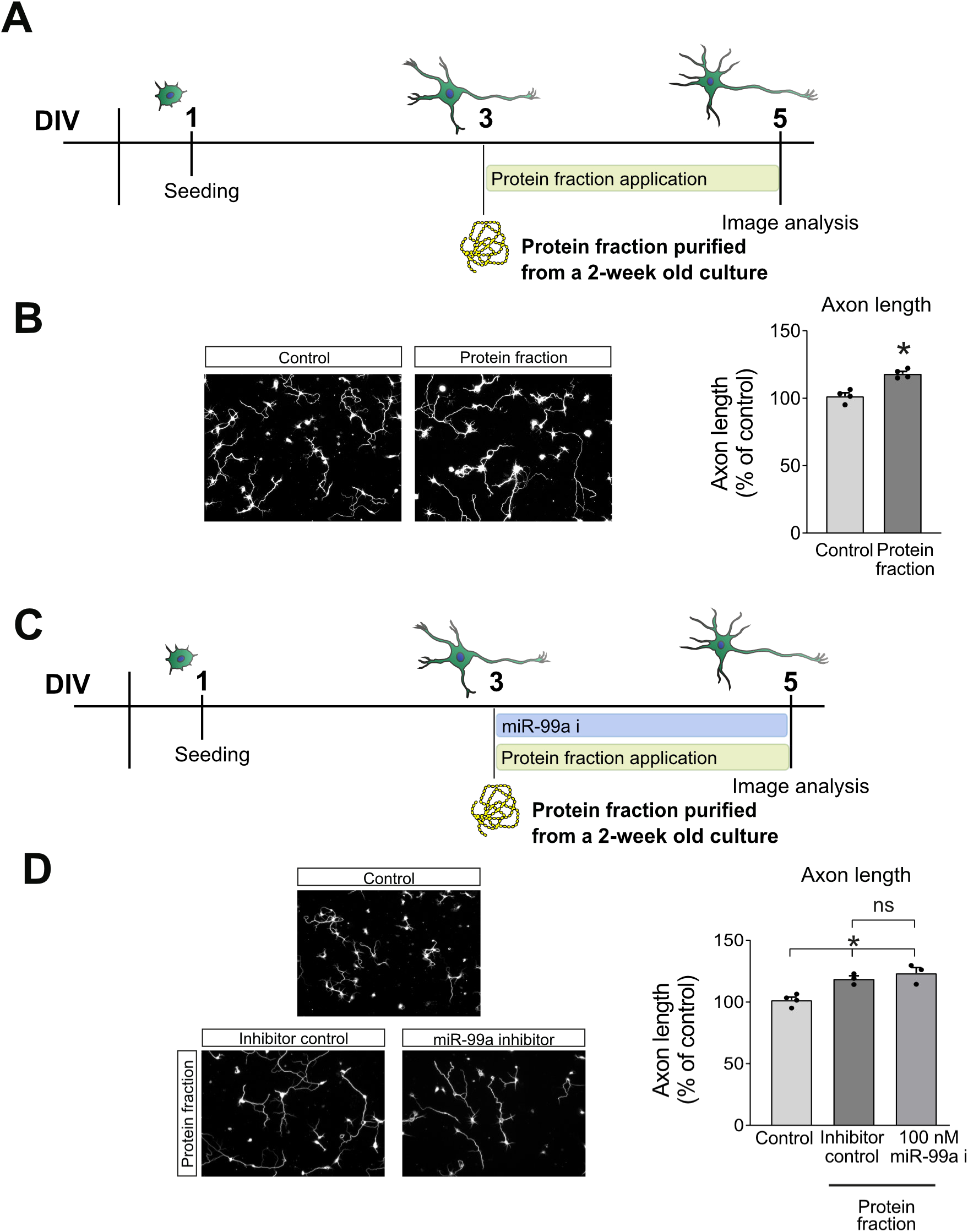
SEC protein-rich fractions from conditioned neuronal media promote modest axonal growth independently of miR-99a. **(A)** Diagrammatic representation of the treatment of DIV3 neurons with protein-rich fractions obtained from neuron culture media by size exclusion chromatography. **(B)** Protein-rich fractions led to a modest increase in axon length in DIV3 neurons after 48h incubation (n=4). **(C)** Diagrammatic representation of the rescue experiment using a cell permeable miR-99a inhibitor alongside the protein-rich fractions. **(D)** The effect of the protein-rich fractions was not rescued after inhibition of miR-99a which suggest the effect on axonal growth is likely associated with the high concentration of growth factors present in neuronal media and eluting in the protein fractions (n=3). All data presented as percentage of control, mean±SEM, * p≤0.05.

**Supplementary Figure 4.**
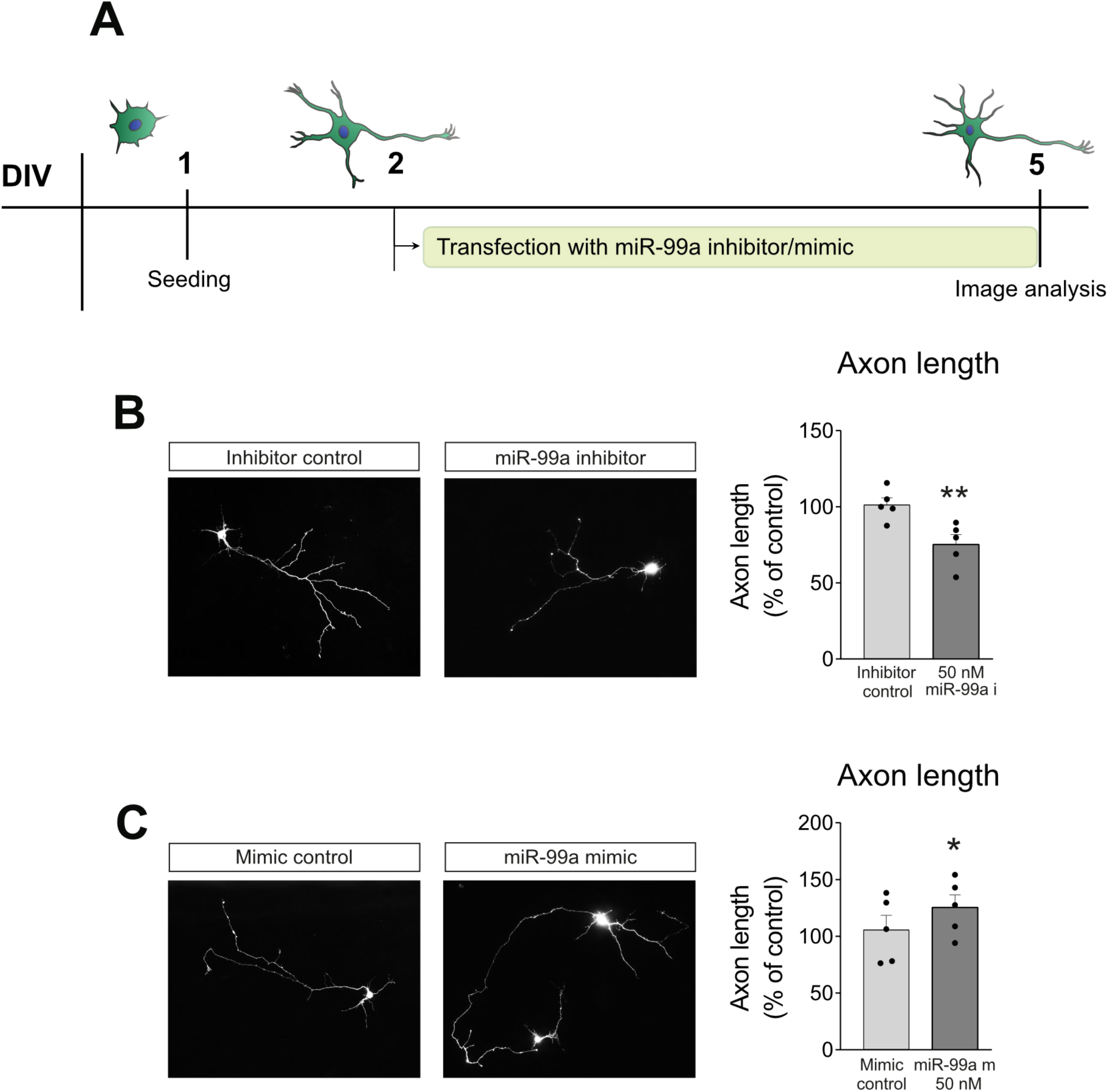
miR-99a regulates axonal outgrowth in cortical primary neurons. **(A)** Diagrammatic representation of co-transfection of DIV2 cortical primary neurons with miR-99a specific inhibitor and mimic oligonucleotides and pGFP (a transfection reporter) for 72h before fixation and imaging. **(B)** Analysis of axon length demonstrated that inhibition of miR-99a leads to a reduction in axonal outgrowth (n=5), **(C)** whereas gain of function experiments using a miR-99a mimic showed an increase in axon length 72h after transfection (n=5). All data presented as percentage of control, mean±SEM, * p≤0.05 and ** p≤0.01.

